# Cilia SubQ, a modular suite of pipelines for automated analysis of primary cilia and ciliary subdomains

**DOI:** 10.64898/2026.04.10.717631

**Authors:** Elizabeth Menzel, Karama Hamdi, Gracie Hoffman, Abdelhalim Loukil

## Abstract

Primary cilia are conserved, antenna-like organelles that protrude from the surface of most vertebrate cells. They function as specialized sensory compartments that detect extracellular cues and convert them into downstream signaling events essential for embryogenesis and tissue homeostasis. The cilium is physically separated from the cytoplasm and contains specialized subcompartments, including the basal body, the transition zone, and the ciliary tip, all of which are critical for its structure, function, and signaling output. Mutations in ciliary genes that disrupt these subcompartments can lead to a wide range of developmental disorders collectively known as ciliopathies. Analysis of these submicron ciliary domains is often time-consuming, repetitive, and prone to user bias. In addition, automated tools for subdomain analysis remain limited, requiring the development of novel, unbiased, and precise segmentation tools applicable to both healthy and pathological conditions. Here, we introduce Cilia SubQ, a versatile suite of flexible pipelines for ZEISS arivis Pro that enables segmentation of the primary cilium, pericentriolar material, basal body, transition zone, and ciliary tip, achieving an approximately eightfold reduction in analysis time with controlled manual intervention. These pipelines are built on our newly developed Cilia.AI, a machine-learning model that recognizes primary cilia with minimal manual correction. The suite also includes a validated script for generating kymographs to assess intraflagellar transport (IFT) dynamics in mammalian primary cilia. Cilia SubQ files and video tutorials are publicly available through the Open Science Framework (OSF) at https://osf.io/hm38f/. Together, the Cilia SubQ pipelines provide batch, high-throughput, and reproducible quantification of primary cilia and ciliary subdomains, delivering greater data output with reduced hand-on analysis.

## Introduction

From early embryogenesis through adulthood, most vertebrate cells harbor a single hair-like organelle called the primary cilium that acts as a cellular sensory antenna, essential for embryonic development and for tissue homeostasis^1,2^. As a specialized signaling hub, the cilium is enriched with a diverse repertoire of molecules, including Sonic Hedgehog (SHH) pathway components, G protein–coupled receptors (GPCRs), and effectors that shape local second-messenger dynamics (e.g., cAMP, Ca²⁺). Consequently, it can detect a variety of extracellular cues and tune multiple downstream signaling pathways^2–7^. Structurally, the cilium consists of a microtubule-based core, the axoneme, encased within the ciliary membrane. Along the core, ciliary trafficking is primarily driven by kinesin and dynein motors that move anterogradely from the ciliary base to the tip and back. These carry the primary transport machinery, Intraflagellar Transport (IFT) complexes and their cargoes, which mediates cilium assembly, structural integrity, and signaling. The BBSome is a specialized cargo adaptor complex that, in concert with IFT, mediates the transport of signaling molecules, particularly their exit from cilia^8–12^.

Ciliary regulation and function are intricately tied to proteostasis within the cilium and at the centrosome, the cell’s microtubule-organizing hub. The centrosome is composed of a mother and a daughter centriole (DC) connected by fibrous linkers, which are embedded within an amorphous proteinaceous matrix called the pericentriolar material^13–16^. During ciliogenesis, the mother centriole undergoes key molecular transitions and is converted into a basal body from which emanates the axoneme. Completion of basal body maturation and initiation of cilium assembly require the recruitment of tau tubulin kinase 2 (TTBK2), removal of the inhibitory capping protein CP110, and engagement of the intraflagellar transport (IFT) machinery^17^. After assembly, spatiotemporal regulation of protein composition is necessary to preserve ciliary architecture and tune signaling. This relies on orderly molecular trafficking that promotes selective entry and retention of specific proteins, or their exit from the cilium in response to specific extracellular stimuli^18,19^. Accordingly, mutations that disrupt ciliary structure or trafficking can alter this equilibrium and cause rare developmental disorders collectively termed “ciliopathies’’^20–23^.

At the ciliary base, a highly organized region termed the transition zone (TZ), also referred to as the ciliary barrier, mediates molecular selectivity and trafficking. The transition zone is characterized by microtubule-ciliary membrane connectors, or Y-links, which are organized by conserved protein modules, including the MKS, NPHP, and CEP290 modules^24,25^. In addition, the ciliary necklace forms a specialized membrane domain surrounding the axonemal base. Mutations in TZ genes can impair the entry and/or exit of signaling molecules, altering ciliary composition, structure, and signaling^26–29^. Another key ciliary subdomain is the ciliary tip (CT), which plays a central role in ciliary function. During SHH activation, the tip accumulates several SHH regulators, including GLI transcription factors and the atypical kinesin KIF7^30,31^. Proper regulation of GLI proteins at the ciliary tip is essential for SHH activation^32^. The same subdomain also serves as a site for intraflagellar transport turnaround, where anterograde trains are remodeled into retrograde trains^33–35^.

The small size of primary cilia, typically a few microns in length, and the submicron dimensions of the basal body make manual measurements laborious and susceptible to bias. Quantification requires counting cilia and measuring pixel-based fluorescence values across multiple fields of view, images, and experimental conditions, rendering the analysis time consuming and repetitive. The challenge is amplified when ciliary-centrosomal subcompartments need to be evaluated, including the basal body, the transition zone, or the ciliary tip. Under fluorescence imaging, these structures generally fall within the submicron range, with apparent dimensions spanning from a few hundred nanometers to <1 µm, depending on the marker, microscopy modality, image-processing and segmentation setting^36–38^. A number of tools have been produced to automate primary cilia annotation and quantify ciliary morphology, length, and fluorescence in mammalian cells^39–43^. While these workflows have advanced the field, automated analysis of ciliary subdomains remains incomplete. There is therefore a need for a modular, high-throughput platform that integrates cilium detection with ciliary subdomain analysis in both healthy and disease states.

Here, we present Cilia-SubQ, a versatile suite of flexible pipelines, which streamlines the detection and quantification of primary cilia (ARL13B+) and key ciliary subdomains, including the basal body, transition zone, and ciliary tip. Using the ZEISS arivis Cloud AI toolkit, we developed Cilia.AI, a model that segments primary cilia while reducing the need for manual correction, enabling additional pipeline steps for the selection and quantification of individual ciliary subdomains. We also provide a validated script for generating kymographs of IFT trafficking in mammalian primary cilia. In this manuscript, we focus on practical application and provide detailed instructions for adapting the pipelines to specific analytical needs. The Cilia SubQ suite files and video tutorials are available through the Open Science Framework (OSF) repository at https://osf.io/hm38f/.

## Materials and methods

### Cell Culture

Wild-type and two independent CRISPR-engineered Csnk2a1 knockout mouse embryonic fibroblast (MEF) clones were cultured in high-glucose Gibco Dulbecco’s modified Eagle medium (DMEM) supplemented with 10% fetal bovine serum (FBS)^44^. Retinal pigment epithelial cells (RPE-1) were cultured in Dulbecco’s Modified Eagle Medium/Nutrient Mixture F-12 (DMEM/F-12) supplemented with 10% FBS. Human skin fibroblasts were cultured in high-glucose DMEM containing sodium pyruvate and GlutaMAX, supplemented with 10% FBS. All cells were maintained in a humidified incubator with 5% CO2 at 37°C. To induce ciliogenesis, MEFs and human skin fibroblasts were serum-starved in their respective base media containing 0.5% FBS for 24–48 h. RPE-1 cells were serum-starved in DMEM/F-12 without FBS for 24–48 h.

### Antibodies

Primary antibodies used for IF were as follows: anti-AHI1 (IF: 1:250, Proteintech, 22045-1-AP), anti-ARL13B (IF: 1:1000–1:1500, Abcam, ab136648), anti-Centrin2 (IF: 1:1000, Sigma, 04-1624), anti-γ-tubulin (IF: 1:1000–1:1500, Sigma, T6557.2ML), anti-GPR161 (IF: 1:200, Proteintech, 29328-1-AP), anti-IFT81 (IF: 1:100, Proteintech, 11744-1-AP), anti-KIF7 (IF: 1:200, Proteintech, 24693-1-AP), anti-polyglutamylated tubulin (IF: 1:500, Adipogen, AG-20B-0020-C100), and anti-TMEM67 (IF: 1:200, Proteintech, 13975-1-AP).

### Immunofluorescence

Cells were plated on 12 mm glass coverslips and grown to approximately 80% confluence. In some experiments, coverslips were pre-treated with gelatin for 45 min before cell plating. Once cells reached confluence, they were serum-starved to induce ciliogenesis. Coverslips were fixed either in 4% paraformaldehyde (PFA) for 5–20 min or in 100% methanol at –20°C for 3–5 min. PFA-fixed cells were subsequently permeabilized in 0.4% Triton X-100 for 10 min.

Fixed cells were blocked in blocking buffer (2% FBS and 0.1% Triton X-100 in PBS) for 1–3.5 h at room temperature. Primary antibodies were incubated overnight (16–20 h) at 4°C with rocking, followed by incubation with secondary antibodies for 1–2 h at room temperature with rocking (1:500; Alexa Fluor 488, Alexa Fluor 568, and Alexa Fluor 647; Molecular Probes, Thermo Fisher Scientific). DNA was stained with DAPI for 3 min. Coverslips were mounted onto glass slides with ProLong Gold mounting medium and allowed to polymerize overnight before imaging.

### Imaging

Cells were imaged using one of the following systems: (1) a Zeiss Axio Observer-Z1 widefield microscope equipped with a Zeiss Plan-Apochromat 63×/1.4 oil objective, an Axiocam 506 monochrome camera, and Apotome optical sectioning; (2) a Nikon Eclipse Ni-E widefield microscope equipped with a Nikon Plan-Apo 60×/1.40 oil objective and an ORCA-Flash4.0 digital camera; or (3) a Nikon Eclipse Ti-2 confocal microscope equipped with a Nikon Plan-Apo 60×/1.42 oil objective and an ORCA-Fusion BT digital camera. All fields of view were acquired as Z-stacks and processed in ImageJ into 8-bit maximum intensity projections for standardized downstream 2D analysis within the SubQ workflow.

### Datasets used for training the Cilia.AI model

We used Zeiss arivis Cloud to train the cilia annotator model on 8-bit maximum intensity projections generated from multiple cell and tissue datasets selected to capture a broad range of ciliary morphologies and signal patterns. Baseline control datasets included untreated wild-type MEFs, untreated RPE-1 cells, and two wild-type human skin fibroblast lines. To represent short cilia, we included four human skin fibroblast lines carrying mutant CSNK2A1 and RPE-1 cells transfected with control, CCT2, or CCT5 siRNAs using Oligofectamine (Invitrogen), followed by recovery and serum starvation. To represent long cilia, we included two untreated Csnk2a1 knockout MEF clones and RPE-1 cells treated with 3 µM SGC-CK2-1, a CK2 inhibitor^45^, prior to fixation, with or without 200 nM SAG.

To capture SHH signaling– and trafficking-related ciliary changes, we included wild-type MEFs, RPE-1 cells, and wild-type human skin fibroblasts treated with 200 or 400 nM SAG during serum starvation for 24–48 h. We also included RPE-1 cells treated during serum starvation with 3 µM SGC-CK2-1N, an inactive structural analog and recommended negative control for the CK2 inhibitor SGC-CK2-1^45^, with or without 200 nM SAG, to provide additional control for pathway-perturbed conditions. To further diversify the training dataset beyond monolayer cell culture, we included immunostained cortical and midbrain sections collected at P31–P32 from FVB Arl13b-mCherry transgenic mice treated with AAV-BirA2 or AAV-MCHR1-BirA2. Expression of these recombinant proteins was not associated with a significant alteration in cilia length under our experimental conditions^46^.

### Datasets used for Validation of Cilia SubQ pipelines

Cilia.AI was validated using previously unseen images from untreated wild-type MEFs, SGC-CK2-1N-treated RPE-1 cells, and untreated wild-type human skin fibroblasts. SubQ_BB_DC was validated using these same datasets together with an additional previously unseen dataset of untreated RPE-1 cells. SubQ_CT was validated using previously unseen datasets from control siRNA-treated RPE-1 cells and untreated wild-type human skin fibroblasts, whereas SubQ_TZ was validated using previously unseen datasets from untreated wild-type MEFs and untreated wild-type human skin fibroblasts. Cilia.AI performance in tissue sections was further assessed using previously unseen cortical and hippocampal brain images generated from the mouse cohort described above. In addition, performance was also examined on training datasets described above to compare behavior between training and non-training datasets.

### Live imaging of primary cilia trafficking

Mouse embryonic fibroblasts (MEFs) stably expressing CSNK2A1-WT-GFP, used as a control condition, were maintained in DMEM supplemented with 10% fetal bovine serum (FBS) and Geneticin (500 µg/mL). For routine culture, cells were thawed and plated in 10-cm dishes, incubated at 37°C with 5% CO2, and the medium was replaced the following day with fresh selective medium. Cells were then seeded into ibidi 8-well slides at 70–80% confluency and incubated overnight at 37°C. The following day, cells were transfected using Lipofectamine 2000 according to the manufacturer’s instructions. For two small wells, 0.7 µg IFT88-GFP and 0.3 µg Arl13b-mCherry were mixed in 100 µL Opti-MEM and combined with 100 µL Opti-MEM containing 5 µL Lipofectamine 2000. A total of 100 µL of the transfection mixture was added to each well containing 200 µL medium and incubated for 1 h 15 min, after which the transfection solution was replaced with fresh medium containing 10% serum and cells were incubated overnight. Cells were then serum starved in DMEM containing 0.5% FBS for 48 h to induce ciliogenesis. Live imaging of IFT88 trafficking was performed using a Hamamatsu ORCA-Fusion BT camera mounted on a Nikon Ti2 microscope with a 60× oil-immersion objective (NA 1.42). Short videos were acquired at one frame every 250 ms for 30 s.

### Immunohistochemistry

P31–P32 mice were used, and brain fixation, sucrose cryoprotection, OCT embedding, sectioning, immunofluorescence staining, and imaging were performed as previously described by Loukil et al., 2025 for mouse brain sections^46^.

### Cilia length script

A script was written in FIJI to skeletonize and segment masked images exported from Arivis Pro. This script utilizes the “Skeletonize” plugin currently provided in FIJI^47,48^. Our script opens files from a chosen directory, skeletonizes them, and generates ROIs, color-coded images, and skeleton statistics. Our script then automatically saves the ROIs, color-coded images, and skeleton statistics to a user-defined directory.

### Statistical Analysis

Data are reported as arithmetic means ± SEM. All graphs and statistical analyses were generated and performed using GraphPad Prism 9. Statistical analyses of the percentage of cilia and percentage of selections per image were performed using a one-sample Wilcoxon signed-rank test, with 100% as the hypothetical median. For the number of cilia and number of selections per image, we used the Friedman test for comparisons among three data categories or the Wilcoxon matched-pairs signed-rank test for comparisons between two paired data categories. Average intensity and size measurements were analyzed using unpaired t tests, except for the average GPR161 intensity dataset, which was analyzed using a two-way ANOVA with Tukey’s multiple-comparisons test.

## Results

### Developing and optimizing Cilia.AI module for precise identification of primary cilia

We first sought to develop a specialized artificial intelligence model, Cilia.AI, designed as a foundational platform that can be integrated into adaptable pipelines for both full-cilium and specific ciliary subdomain-level annotation. We selected the ZEISS arivis Pro platform and its associated artificial intelligence training toolkit, the ZEISS Arivis Cloud, to leverage a standardized and flexible modular environment capable of batch analysis across diverse datasets for high-throughput primary cilia quantification. A key challenge we faced was the intrinsic diversity in primary cilium number and morphology across two-dimensional (2D) images from various cell lines and tissues **(Figure 1A)**. To develop Cilia.AI, we first trained the model using 2D images from two widely used cell lines in ciliary research: wild-type mouse embryonic fibroblasts (MEFs) and retinal pigment epithelial (RPE-1) cells. Cells were cultured and plated on coverslips, and then serum-starved to induce ciliogenesis. Immunofluorescence staining was performed on fixed cells using antibodies to ARL13B and γ-tubulin to label cilia and centrosomes, respectively. These markers were intentionally selected because of the availability of well-validated, commercially available reagents and their widespread use in the cilia research community. Widefield microscopy was performed, and z-stack images were processed as maximum intensity projections (MIPs) for analysis.

**Figure 1:**
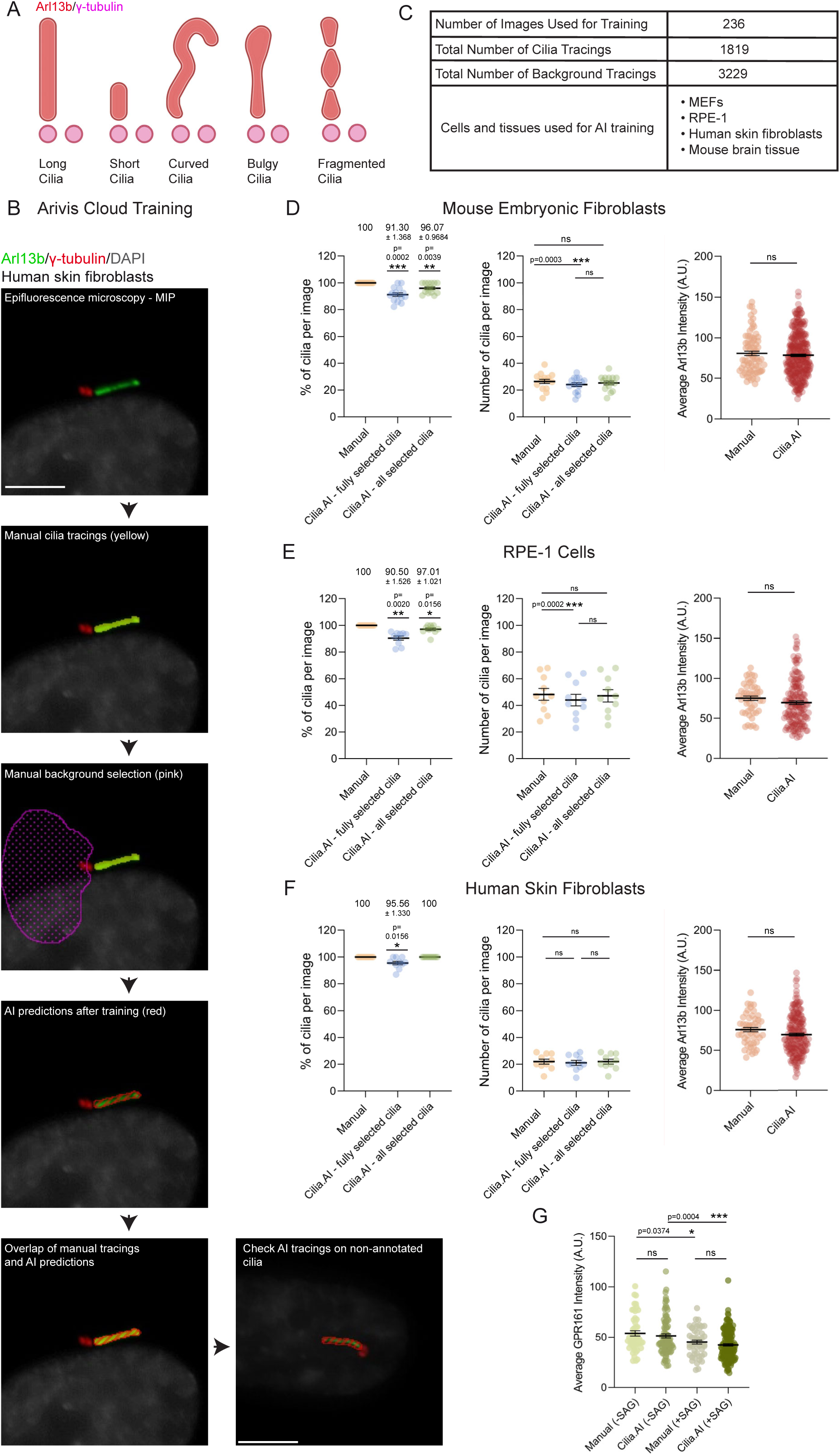
Training and validating Cilia.AI. **A.** Depiction of diverse primary cilia morphologies. **B.** Visualization of how Cilia.AI was trained in Arivis cloud. 8-bit MIP images are imported into Arivis Cloud, and the primary cilium (yellow) and background (pink) are manually traced. The model trains for 24 hours, then the Cilia.AI tracings are compared to the manual tracings and cilia that were not manually annotated (Scale bar: 2 µm). **C.** An overview of the datasets and their contents used to train Cilia.AI v32. **D.** Graphs showing the detection performance of Cilia.AI compared to manual annotations in MEFs. Left graph: the percentage of cilia detected per image, analyzed with a one-sample Wilcoxon signed-rank test. Middle graph: the number of cilia detected per image, analyzed using the Friedman test. Right graph: the average ciliary ARL13B intensity, analyzed with an unpaired t-test (Manual: n=50 cells; Cilia.AI: n=326 cells). **E.** Same as D, but for RPE-1 cells. (Manual: n=50 cells; Cilia.AI n=156 cells). **F.** Same as D, but for human skin fibroblasts. (Manual: n=50 cells; Cilia.AI: n=198 cells). **G.** Graph showing the average ciliary GPR161 intensity in non-SAG-treated cilia detected either manually (n=50 cells) or by Cilia.AI (n=88 cells), and in SAG-treated cilia detected either manually (n=50 cells) or by Cilia.AI (n=149 cells). Statistical analysis was performed using a two-way ANOVA. For panels D-G, a total of nine independent experiments were analyzed (three for D, and two each for E-G). For graphs showing the number and percentage of cilia, a total of 1,099 cilia were quantified across all three cell lines.

Cilia.AI training was conducted using expert manual annotations of ARL13B-positive cilia, including the annotation of non-ciliary fluorescent areas (**Figure 1B)**. During the validation process, the newly trained Cilia.AI model demonstrated variable performance: it successfully detected cilia in MEFs but showed reduced accuracy in RPE-1 cells, which typically exhibit higher ciliary density. Performance was further compromised when analyzing cilia with abnormal morphology or low ARL13B fluorescence, highlighting specific limitations that required further training with additional optimization to enhance model capabilities. We therefore focused on expanding the training dataset by annotating a greater number of images from wild-type MEFs, RPE-1 cells, and human skin fibroblasts. In addition, we exposed Cilia.AI to abnormally formed cilia through a series of ciliary manipulations that alter ciliary length (shortened cilia: siRNA targeting CCT proteins^49^; elongated cilia: Csnk2a1 KO cells^44^) and/or activate the Sonic Hedgehog pathway. Successive Cilia.AI training on these diverse datasets, including those capturing a wide range of ciliary morphologies and ARL13B fluorescent intensities, resulted in improved detection performance.

One round of training within the ZEISS arivis Cloud takes approximately 48 hours to complete, including critical corrective manual annotations. Therefore, at least two separate models were generated per round: one prior to and one after manual corrections were made. Additionally, some model versions required a second round of manual corrections to perform adequately before more images were added for training. To date, we have performed a total of 35 rounds of training, reaching 32 versions of Cilia.AI. In total, we annotated at least 236 images and traced 1,819 primary cilia and 3,229 background tracings **(Figure 1C)**. In this manuscript, we evaluated the performance of our latest version, Cilia.AI v32 **(File name: Cilia.AI v32.czann)**.

To assess the performance of Cilia.AI, we extensively compared its detection accuracy to manual detection performed by an experienced human annotator. Importantly, the testing focused on previously unseen datasets, distinct from those used for training, to assess detection robustness and minimize overfitting. The Cilia.AI model was imported into the ZEISS arivis Pro software using the Deep Learning Segmenter function. During validation, we observed occasional non-ciliary annotations, typically only a few pixels in size, that were incorrectly classified as primary cilia. To address this and limit manual corrections, we incorporated two filtering steps after AI segmentation. The first was an object-feature filter that excluded any Cilia.AI selections below a defined size threshold, chosen so as not to affect the detection of short cilia. The size threshold can be adjusted or removed as needed, but we recommend its use to eliminate non-ciliary selections detected by Cilia.AI. The second was a feedback step that restricted primary cilia selection to objects in proximity to the basal body. Validation was performed across three cell lines to assess accuracy. With the addition of these two filter steps, the AI model consistently achieved an average detection success rate of 90% or higher across all three cell lines (MEFs, RPE-1, and skin fibroblasts) prior to manual correction. This performance was even more comparable (approximately 97% or higher) to manual quantification when partially detected cilia requiring minimal corrections were also counted **(Figure 1D, E, and F; left and middle graphs)**. Notably, the detection performance was similar on the training datasets, suggesting that the model has likely reached a performance plateau **(Figure S1A)**. Although manual corrections were minimal under our conditions, as shown by the number of cilia per image, we still recommend visual confirmation of the AI annotations to ensure precision. Following Cilia.AI segmentation, we confirmed that ARL13B fluorescent intensity measurements did not statistically differ from manual quantifications across all three cell lines **(Figure 1D, E, and F; right graphs)**. It is worth noting that additional outputs such as sum intensity and projected area are also available through the ZEISS arivis Pro software.

To further validate the Cilia.AI model, we tested whether cilia selections generated from Cilia.AI-based ARL13B staining could be used to quantify another ciliary marker. Specifically, we focused on primary cilium signaling and activation of the SHH signaling pathway. In the inactive state, SMO, a positive regulator of SHH, is prevented from entering the cilium by PTCH1. In contrast, primary cilia harbor the negative regulator of SHH, GPR161. In the presence of the SMO agonist, SAG, SMO accumulates and GPR161 exits the cilium, leading to discernible changes in protein levels that provide a robust readout for quantification^50,51^. We treated wild-type skin fibroblasts with SAG to assess changes in ciliary GPR161 levels, comparing manual quantification with Cilia.AI–based analysis **(Figure 1G)**. Both approaches showed that SAG treatment resulted in reduced ciliary GPR161 levels compared with untreated cells. Importantly, there was no significant difference between the two analysis methods in either the absence or presence of SAG. This indicates that Cilia.AI–based annotations can be used to reliably assess changes in protein levels associated with primary cilia signaling.

During the training process of Cilia.AI, we also included a set of training sessions to recognize primary cilia in immunostained sections from mature mouse brain. Primary cilia in the brain display a wide range of morphological diversity, providing ideal conditions to enhance model recognition. Sagittal brain sections were prepared from P31–32 Arl13b-mCherry transgenic mice. Brain slices were stained with ARL13B antibodies and DAPI. Although the training dataset included only a limited number of images from cortical and midbrain regions, Cilia.AI, trained primarily on cell-based images, was able to detect an average of 69 ± 3.3% of primary cilia in mature cortical and hippocampal brain sections. This average increased to 87.1% ± 3.78% when partially selected cilia were included, requiring manual corrections. Only about 13% were not detected by Cilia.AI, primarily because of lower Arl13b intensity comparable to non-ciliary signal **(Figure S1B, C)**. These results suggest the potential versatility of this model in detecting primary cilia in a more challenging in vivo context.

We then compared cilia detection time between Cilia.AI, including corrections, and fully manual quantification. Cilia.AI reduced the total analysis time from image import to data export by approximatively 7.5-fold **(Figure S1D)**. Altogether, Cilia.AI-driven analysis significantly reduces hands-on time and increases data output, making it an effective tool for ciliary quantification.

### Segmentation and analysis of the basal body and daughter centriole

We then sought to build a pipeline capable of distinguishing the basal body from the daughter centriole, streamlining the analysis of centrosomal proteins. To this end, Cilia.AI annotations were taken as the reference point to identify the first and second nearest centriolar objects. We first labeled wild-type MEFs with γ-tubulin, a pericentriolar material (PCM) marker, aiming to use its broad centriolar selection as the focal point for annotating both centrioles. A blob finder function was applied to broadly detect small, uniform puncta based on user-defined parameters (for one experiment in MEFs: a split sensitivity of 100%, a diameter of 4 pixels, and a probability threshold of 21.94%) **(Figure 2A)**. The latter two parameters must be adjusted for each centrosomal marker and each independent dataset, while the split-sensitivity threshold must remain at 100% to ensure accurate separation of individual centrioles. The basal body was identified as the centrosomal nearest object to the primary cilium, based on computed cilium–centriole distances. Our findings showed that the generated pipeline, SubQ_BB_DC, identified an average of 92.61% ± 1.47% of correct basal bodies with no corrections **(Figure 2B, SubQ_BB_DC.zpipeline, Video S1)**. The remaining 7.39% consisted of cases where either both centrioles were detected as a single object, or the daughter centriole was incorrectly identified as the basal body and vice versa. Minimal corrections can be applied to these instances by re-selecting the correct objects and assigning the appropriate basal body tag. Incorrectly labeled “basal body” selections should also be deleted.

**Figure 2:**
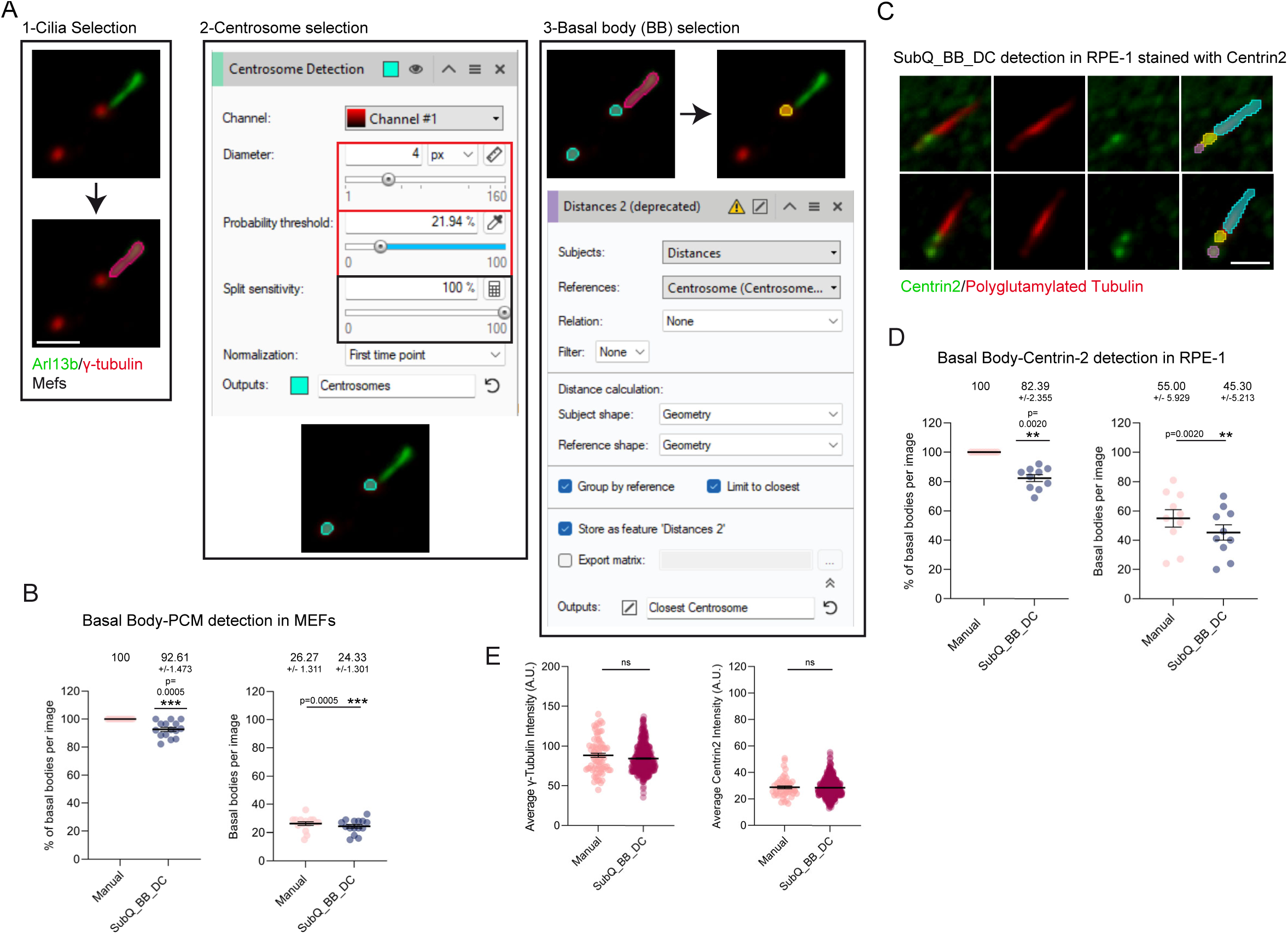
Constructing and validating a pipeline for basal body detection. **A.** Screenshots illustrating key steps of the SubQ_BB_DC pipeline in the Zeiss Arivis Pro used to detect the basal body in MEFs labeled with ARL13B (green) and γ-tubulin (red); scale bar: 2 µm. **B.** Graphs showing the basal body detection performance of SubQ_BB_DC in MEFs. The left graph shows the percentage of basal bodies correctly detected per image, analyzed using a one-sample Wilcoxon signed-rank test. The right graph shows the number of basal bodies detected per image analyzed using the paired Wilcoxon Signed-Rank Test. **C.** Representative images of primary cilia in RPE-1 cells stained with polyglutamylated tubulin (red) and Centrin-2 (green). The overlays depict SubQ_BB_DC detection of the basal body in yellow, and daughter centriole in pink on cilia. Polyglutamylated tubulin selections were traced manually in blue (scale bar: 2 µm). **D.** Same as B, but for RPE-1 cells co-labeled with polyglutamylated tubulin and Centrin-2 **E.** Left graph showing the average γ-tubulin intensity of basal bodies in MEFs detected either manually (n=75 cells) or with SubQ_BB_DC (n=393 cells). Right graph showing the average Centrin-2 intensity in RPE-1 basal bodies detected either manually (n=50 cells) or with SubQ_BB_DC (n=480 cells). Statistical analysis was performed for both using an unpaired t-test. For panels B, D, and E, data were collected and analyzed from a total of four independent experiments. For panels B and D, a total of 944 cilia were quantified across both cell lines.

Unlike MEFs, RPE-1 cells and skin fibroblasts displayed more closely apposed centrioles. We therefore asked whether γ-tubulin could be used to distinguish between individual centrioles. The SubQ_BB_DC pipeline was therefore tested on both cell lines co-stained with ARL13B and γ-tubulin. Our testing indicated reduced detection compared with the pipeline’s performance seen in MEFs, even after adjusting diameter and probability threshold parameters **(Figure S2)**. The percentage of incorrectly detected basal bodies was 45.38% for human skin fibroblast and 56.32% for RPE-1 cells **(Figure S2A and B)**, showing the limitation of the γ-tubulin marker for resolving closely joint centrioles, which often appeared as a single PCM blob **(Figure S3A)**. Changing the selection diameter and probability threshold was also insufficient to properly identify the two centrioles **(Figure S3B)**. To address this challenge, we labeled RPE-1 cells with antibodies to polyglutamylated tubulin and Centrin-2, a core centriolar marker, which allowed for a more defined labeling of centrioles. As Centrin-2 structures differed from those of γ-tubulin, the SubQ_BB_DC pipeline had to be adjusted with a smaller diameter of 2 pixel and a probability threshold to 20.67%, ensuring selection of all Centrin-2-positive signals **(Figure 2C)**. Under these conditions, an average of 82.39% +/− 2.35% of basal bodies were correctly detected with no manual correction, indicating the adaptability of the pipeline to additional centriolar markers in cells with less separated centrioles **(Figure 2D).**

To further validate the SubQ_BB_DC pipeline, we assessed how reliable the automated basal body selection recapitulated the manual method. We evaluated one of the key outputs, average fluorescence intensity, of γ-tubulin– or Centrin-2-labeled selections in MEFs and RPE-1 cells, respectively. We found no significant difference between the two methods in either cell lines, generating high-density datasets with reduced standard error of the mean (SEM) and providing high confidence in the reliability of the analysis **(Figure 2E)**.

We previously discovered that the daughter centriole can facilitate the maturation of the mother centriole into a basal body, driving cilia assembly^16^. The daughter centriole is one generation younger than the mother centriole and can concentrate common centrosomal markers, as well as more specific factors such as Centrobin, Cep120, and Neurl-4^52–54,16^. To expand the capabilities of the centrosome/cilia field for studying this organelle, we extended the SubQ_BB_DC pipeline to specifically annotate the daughter centriole, building on our initial centrosome detection coupled with basal body identification (γ-tubulin in MEFs and Centrin-2 in RPE-1 cells) **(Figure 2)**. To this end, we isolated all objects identified as centrosomes but not tagged as basal bodies. We then applied a secondary distance-based criterion to select the next-closest centrosomal object to the primary cilium, which was categorized as the daughter centriole **(Figure S3C)**. We found that the SubQ_BB_DC pipeline enabled automated identification of 87.1 +/−1.89% of daughter centrioles in MEFs and showed lower performance in RPE-1 cells (75.91% +/− 3.19%) **(Figure S3D, E)**. This was expected, as secondary detection of the daughter centriole heavily relies on the performance of the initial basal body detection.

The final SubQ_BB-DC pipeline, comprised of 11 steps, provides a flexible framework for automated dual annotation of most basal bodies and daughter centrioles, serving as a foundation for researchers to quantify centrosomal protein levels, foci number, projected area, and other morphometric parameters. Pipeline performance is linked to the quality of centrosomal protein staining and accurate cilium detection. A series of manual corrective steps may be required to achieve comprehensive centriole selection across all cells. Overall, the SubQ pipeline effectively reduces labor-intensive, repetitive tasks and facilitates higher-throughput analysis of centrosomal compartments.

**Critical notes:** For basal bodies and daughter centrioles that were incorrectly selected, a series of simple steps can be taken to recover these data points. First, we identify selections tagged with “Basal body (or DC) OUTPUT” that are not basal bodies (or DCs). We either remove their tag or delete these selections. We then find the correct blob finder selections of basal bodies (or DCs) or manually annotate the missing ones. Finally, we add the tag “Basal body (or DC) OUTPUT” to these new objects to be exported with the automated annotations. For centrioles identified as a single object, we recommend deleting it and manually tracing the basal body by assigning the “Basal body OUTPUT” tag to the newly created selection. In this context, a non-daughter centriole will be selected, as there is no distance limit to the second closest object to the primary cilium. This will most likely be a centriole from a neighboring cell and that object should also be deleted. We also recommend removing the single selection, manually tracing the daughter centrioles, and adding the tag “Daughter centriole OUTPUT”.

### Building and validation of the ciliary barrier subdomain

Primary cilia contain a structural barrier called the transition zone, which separates the cilium from the cell cytosol. This compartment selectively regulates the entry and exit of ciliary proteins and is considered a disease hotspot for multiple ciliopathies^55,29,28^. Despite its critical role, the transition zone spans only a few hundred nanometers, making it challenging to analyze. Using the Cilia.AI model, we aimed to build a pipeline that systematically detects a transition zone marker. Wild-type human skin fibroblasts were serum-starved to induce ciliogenesis and then stained with ARL13B, γ-tubulin, and AHI1, a known transition zone marker. Following cilium and basal body detection, we applied another “blob finder” command designed to identify blob-like structures in the AHI1 channel. This step was meant to broadly select foci-like structures and was not expected to distinguish between ciliary and non-ciliary fluorescence signals. For AHI1, we used the following parameters: a diameter of 5 pixels, a probability threshold of 30.32%, and a split sensitivity of 65.23%, which are key variables that must be retuned for each transition zone marker or dataset **(Figure 3A)**. At this point, the annotations included all transition zone signals as well as additional non-ciliary selections. To begin improving specificity, we identified and expanded the nearest PCM blob (by 6 pixels; adjustable) with a “Segment Morphology” function. This variable was refined so that the PCM selection overlapped with the proximal side of the cilium. We then forced the intersection of the dilated PCM with the cilium, selecting the cilium’s proximal extremity and creating a “transition zone area” **(Figure 3A)**. We then retained only AHI1 blob finder results that were touching, overlapping, or within 2 pixels of the transition zone area. All other results were discarded to eliminate most non-specific transition zone selections. As a second failsafe mechanism, we incorporated a distance function to instruct the transition zone pipeline to consider only the closest blob to the basal body. As a result of these layers of stringency, only the puncta associated with the transition zone area and closest to the basal body were included in the final segmentation **(Figure 3B, Figure S4)**. Under these conditions, an average of 99.5% +/− 0.5 of AHI1 transition zone signals was selected with no manual corrections **(Figure 3C, SubQ_TZ.zpipeline, Video S2)**. The SubQ_TZ pipeline lays the groundwork for flexible and reliable workflow that can facilitate the precise and unbiased quantification of TZ proteins.

**Figure 3:**
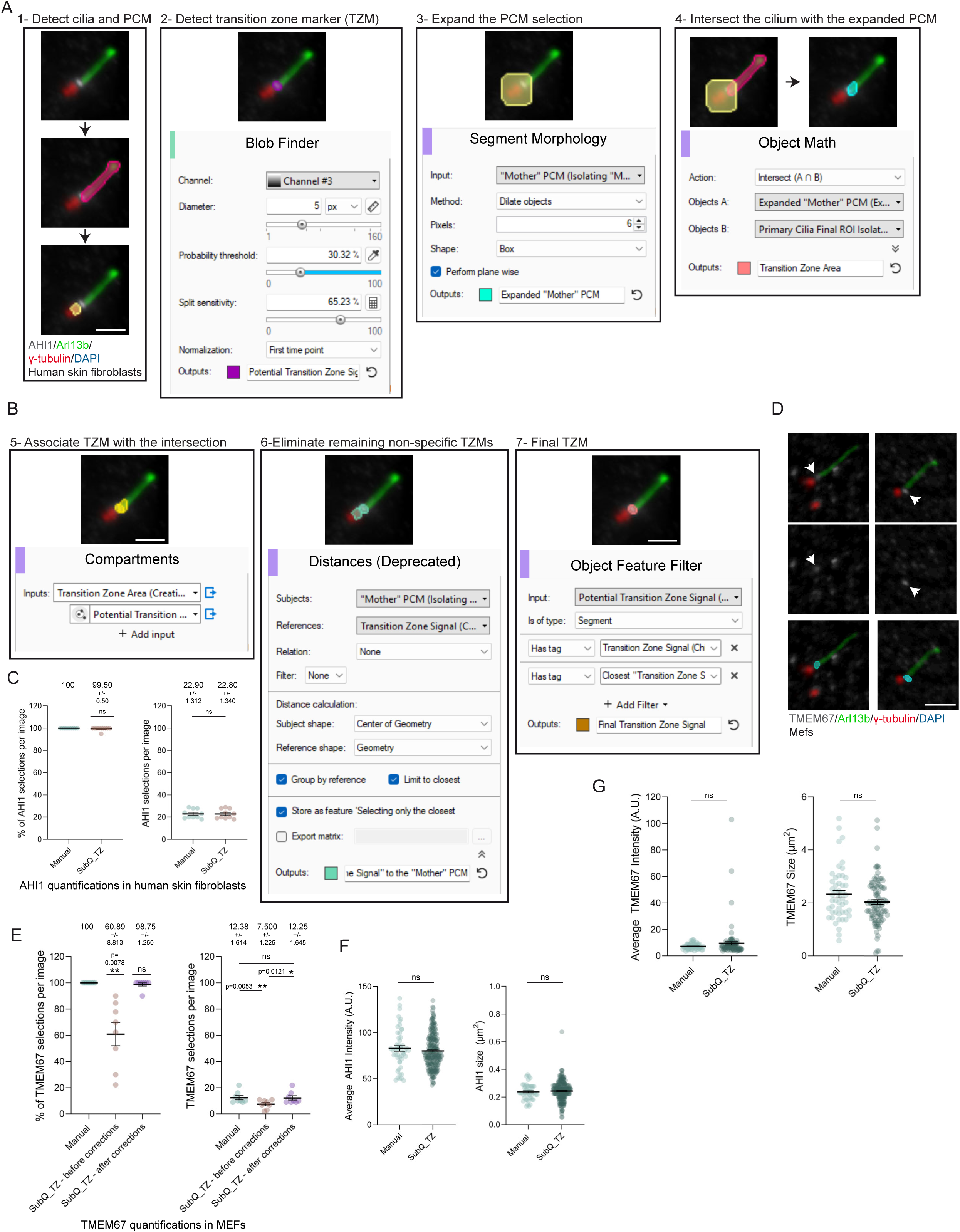
Building a transition zone detection pipeline and testing its performance. **A.** Visualization of the specific SubQ_TZ pipeline operations used to create a “transition zone area,” with an example of each step depicted alongside AHI1-stained human skin fibroblasts (scale bar: 2 µm). **B.** Depiction of the SubQ_TZ pipeline steps used to identify and isolate the correct transition zone signal, with a representative example of AHI1-stained human skin fibroblasts (scale bar: 2 µm). **C.** Graphs displaying the transition zone detection performance of SubQ_TZ in human skin fibroblasts stained with AHI1. Left graph: the percentage of AHI1 transition zone signals correctly detected per image (one-sample Wilcoxon signed-rank test). Right graph: the number of AHI1 transition zone signals correctly detected per image (paired Wilcoxon signed-rank test). **D.** Representative images of MEFs stained for TMEM67 (gray), Arl13b (green), and γ-tubulin (red). SubQ_TZ detection results for TMEM67 transition zone signal overlaid on cilia (cyan, scale bar: 2 µm). **E.** Graphs portraying SubQ_TZ transition zone detection performance in MEFs stained for TMEM67. Left graph: percentage of TMEM67 transition zone signals correctly detected per image (one-sample Wilcoxon signed-rank test). Right graph: number of TMEM67 transition zone correctly detected per image (Friedman’s test). Both include statistics before and after manual corrections. **F.** Graphs showing the average AHI1 intensity (left) and projected area (right) of transition zone signals detected either manually (n=50 cells) or with SubQ_TZ (n=230 cells). Statistical analysis performed using an unpaired t-test. **G.** Graphs showing the average TMEM67 intensity (left) and projected area (right) of manually detected TMEM67 transition zone signals (n=50 cells) and those detected by SubQ_TZ (n=95 cells). Statistical analysis carried out using unpaired t-test. For panels C, and E-G, data were extracted from a total of four independent experiments (two for C and F, two for E and G). For graphs in C and E, a total of 327 primary cilia were quantified.

To further test this pipeline, we co-stained MEFs with another transition zone protein, TMEM67, in combination with ARL13B and γ-tubulin. Our staining conditions for TMEM67 displayed subtle transition zone localization with non-ciliary foci throughout the cell, providing an opportunity to challenge the SubQ_TZ workflow **(Figure 3D)**. For proper detection of TMEM67 signal, we reduced the probability threshold of the blob finder to ensure a comprehensive selection of all transition zone areas. This resulted in a greater number of non-specific selections incorrectly associated with the transition zone. There was also no longer a guarantee that the single closest “transition zone signal” to the basal body was still the correct transition zone signal. To remedy this, we modified the distance function to include the three closest “transition zone blobs” to the basal body, which resulted in a successful detection of 60.89% +/− 8.813% of ciliary transition zones with no corrections **(Figure 3E)**. At this point, the remaining 39.11% of cilia displayed more than one transition zone selection. The minimal corrective action we used was to delete non-specific transition zone signals, which led to an average detection of 98.75% +/− 1.250% **(Figure 3E)**.

The final SubQ_TZ pipeline is comprised of 16 steps that can deliver key outputs, including fluorescence intensity levels (average and sum) and the projected area, considered phenotypic entry-level parameters for the ciliary barrier. We further assessed SubQ_TZ by comparing it to expert manual annotations for AHI1 and TMEM67 staining. Average intensity and projected area did not differ from the manual datasets **(Figure 3F, G)**. This suggests that the pipeline’s performance is suitable for detecting AHI1 and TMEM67 signals under our experimental conditions. SubQ_TZ streamlines the often-time-consuming process of manual annotation, yielding a larger number of data points for robust statistical analysis and consistency. Overall, SubQ_TZ performance is tightly linked to the intrinsic quality of the transition zone labeling and staining conditions. Additional puncta-like noise can hinder the automated detection of the true transition zone structures. Yet we provide streamlined corrective measures that can be deployed to enhance the versatility of this transition zone annotator.

### Recognition and quantification of ciliary tip subdomain

The ciliary tip is another key subdomain located at the distal terminus of the primary cilium. Similar to the transition zone subcompartment, we sought to build a comprehensive tool that systematically annotates ciliary tip markers. We cultured human skin fibroblasts and induced ciliogenesis in the presence of SAG, which promotes accumulation of SHH proteins at the tip, including GLI2 and KIF7. We therefore co-stained cells with antibodies to ARL13B, γ-tubulin, and KIF7. Similar to previous workflows, the first step deployed cilium and pericentriolar material (basal body) recognition as the foundation of the ciliary tip pipeline. We then applied the Blob Finder to identify foci-like structures across the image (diameter: 2 pixels; probability threshold: 21%; split sensitivity: 0% or 68%). These parameters must be adjusted based on the ciliary tip marker, with the critical requirement that tip signals are fully captured, even though non-ciliary signals may be selected. The third step associated each cilium with its tip counterpart in a parent-child relationship and created categories based on the number of tips detected per cilium **(Figure 4A)**. This classification separated cilia into two groups: those with a single KIF7 selection and those with two or more KIF7 foci. This approach preserved those with a single, unambiguous tip marker with no adjacent non-ciliary structures **(Figure 4B)**. In cases with multiple foci within the ciliary vicinity, these were categorized as “cilia with more than one tip.” In this context, non-tip foci tended to localize to the proximal end of the cilium. To eliminate these objects, we applied a distance operation to identify foci closest to the PCM. An object feature-based filtering step was then used to exclude these proximal foci, retaining only KIF7 selections that were not closest to the PCM while still preserving the associated child feature **(Figure 4A).**

**Figure 4:**
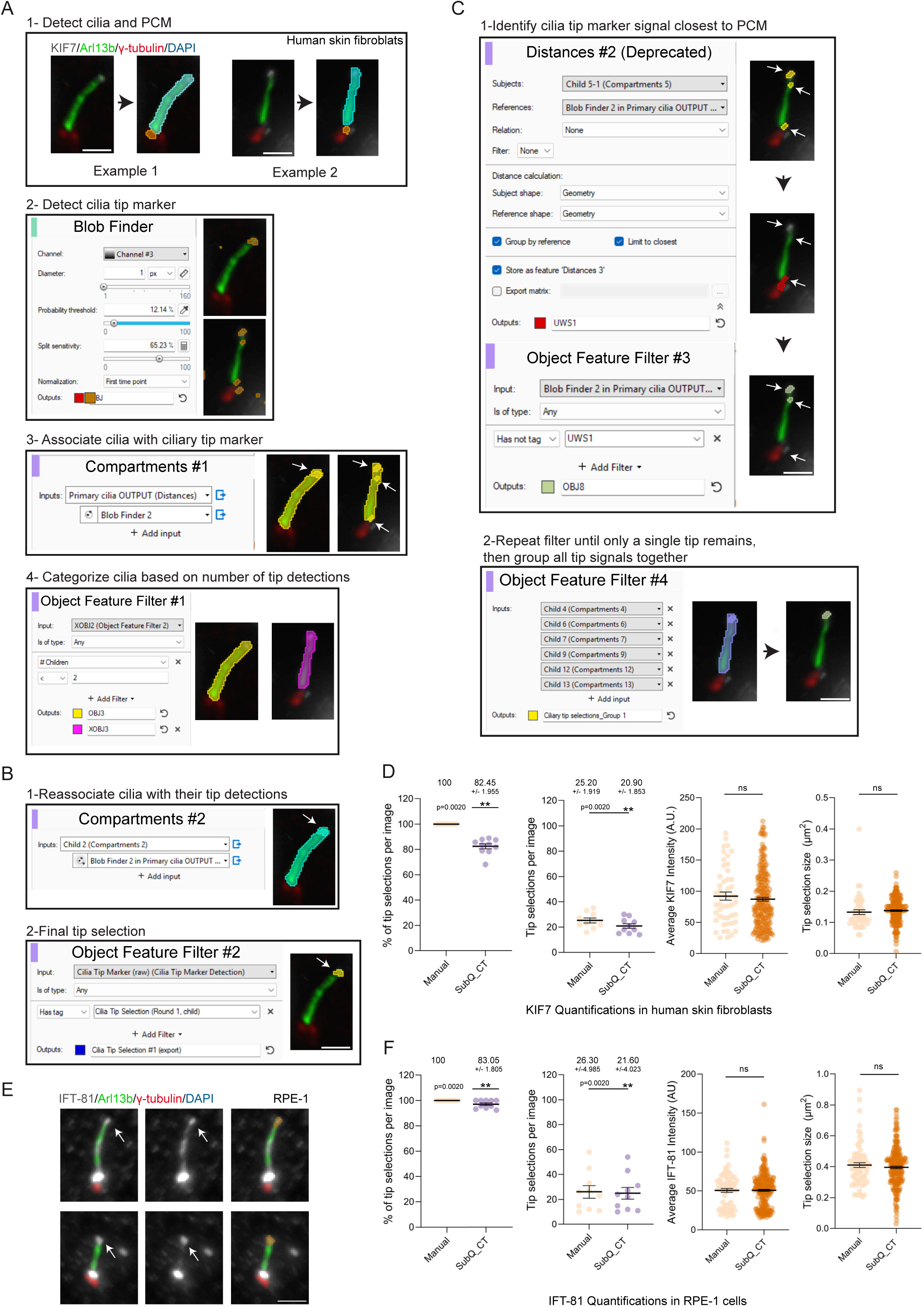
Building and validating a pipeline for ciliary tip detection. **A.** Visualization of the specific pipeline operations implemented in SubQ_CT to initially categorize primary cilia based on the number of signals in proximity to the primary cilium (either one proximal signal or more). The results of each operation are shown for two cilia stained for KIF7 (gray), ARL13B (green), and γ-tubulin (red) in human skin fibroblasts. Cilia, basal body and ciliary tip annotations are shown as colored selections (scale bar: 2 µm). **B.** Depiction of the pipeline steps in SubQ_CT used to isolate the ciliary tip signal from primary cilia with only a single proximal signal detected, along with a representative example of human skin fibroblasts stained for KIF7 (same as A). Cilia and ciliary tip annotations are shown as colored selections (scale bar: 2 µm). **C.** Representation of the pipeline steps in SubQ_CT used to calculate and remove signals based on proximity to the basal body until only a single tip signal remains. A representative example of a cilium stained for KIF7 (same as A) in human skin fibroblasts that had multiple tip signals detected is shown alongside. Cilia and ciliary tip annotations are shown as colored selections (scale bar: 2 µm). **D.** Graphs displaying the ciliary tip detection performance of SubQ_CT in human skin fibroblasts stained with KIF7. The first graph shows the percentage of KIF7 tip selections correctly detected per image, analyzed using a one-sample Wilcoxon signed-rank test. The second graph shows the number of KIF7 tip selections correctly detected per image, analyzed using a paired Wilcoxon signed-rank test. The third and fourth graphs show the average KIF7 tip intensity and projected area, respectively, of manually detected tip signals (n=50 cells) and those detected by SubQ_CT (n=248 cells). Statistical analysis for both was performed using unpaired t-tests. **E.** Representative images of RPE-1 cells stained for IFT81 (gray), Arl13b (green), and γ-tubulin (red). The results of SubQ_CT detection of ciliary tip IFT81 signal are overlaid on cilia (orange, scale bar: 2 µm). **F.** Same as D, but in RPE-1 cells labeled with IFT-81. Manual selections (n=50 cells) and SubQ_CT annotations (n=149 cells). For panels D and F, a total of four independent experiments were analyzed. For the percentage of selections and number of selections per image graphs, a total of 515 cilia were quantified across the two cell lines.

During our pipeline design, we found that repeating the latter step five times was necessary to remove most incorrect ciliary tip annotations and obtain higher accuracy of KIF7 tip selections **(Figure 4B, C, SubQ_CT.zpipeline, and Video S3)**. We then evaluated the performance of the SubQ_CT pipeline in comparison to manual annotation. Our findings showed comparable tip selection per image with an average percentage of successful KIF7 tip selections of 82.45% ±1.955%. Additionally, the average KIF7 intensity and projected 2D area measurements obtained from SubQ_CT and manual annotations were statistically similar, indicating reliable pipeline performance with seamless extraction of key readouts necessary for analysis of the ciliary tip **(Figure 4D)**.

SubQ_CT depends on a robust ciliary tip signal. While SHH pathway regulators such as GLI2 and KIF7 accumulate at the ciliary tip upon pathway activation, tip enrichment is reduced or absent in the absence of SHH agonists, preventing reliable tip selection and leading to potential misidentification of background signal. In addition, ciliary tip levels of these proteins can vary widely between normal and pathological conditions, making such changes difficult to quantify without an independent tip marker. Therefore, relying solely on SHH markers for tip selection is limiting, particularly when these markers are also the targets being measured.

To remedy this, we chose to test SubQ_CT with a more stable ciliary protein, IFT81, predominantly present at the tip and base of the cilium. The goal was to use the IFT81 tip signal as a general scaffold for defining the tip compartment, making it possible to quantify fluorescence signals of other proteins at the tip. This would enhance the ability of the SubQ_CT pipeline to quantify a wider range of ciliary tip proteins and assess their intensity levels in a more unbiased manner. For IFT81 staining, we implemented the following changes to the Blob Finder of IFT81 within the SubQ_CT pipeline: a diameter of 1 pixel and a probability threshold of 12%. While IFT81 localizes to the tip, we observed that a small subset of cilia showed no prominent tip enrichment. To account for these cells, we found that the optimal split sensitivity was 68%, allowing for the segmentation of the IFT81 tip signal in this subset. Our evaluation of the modified SubQ_CT–IFT81 pipeline showed an average tip recognition rate of 83.05% ±1.805%, with performance comparable to manual annotation. The pipeline also yielded similar average values for IFT81 intensity and projected area **(Figure 4E, F)**. The SubQ_CT pipeline comprises 55 steps that can still be optimized to accommodate specific research needs; it also supports the use of alternative tip markers capable of delivering accurate, rapid, and unbiased quantification of the ciliary tip subdomain.

Altogether, SubQ_BB_DC, SubQ_TZ, and SubQ_CT reduced the burden of repetitive manual annotation and provided comparable gains in overall analysis time. Based on timed transition zone annotation, the SubQ pipelines reduced hands-on analysis time by approximatively 8-fold relative to manual quantification **(Figure S4D)**, supporting the utility of the SubQ framework for subdomain-level ciliary analysis.

### Critical notes on priming pipelines, SIS files, and batch analysis

All pipelines require an initial priming step in which the Cilia.AI model is imported before any analysis can proceed **(Cilia.AI v32.czann)**. This operation is executed through the ‘’Deep Learning Segmenter’’ **(Figure S5A)**. We usually assign the centrosome marker to the first position and the cilia marker to the second. If performed correctly, the priming step is completed only once, and the pipeline is exported for future use **(Figure S5B, Video S4)**. We also recommend a couple of checks to ensure the pipeline is running as intended before generating data for analysis. Each step in the pipeline can be run by clicking the arrow in the top left corner of the analysis panel, which will trigger the next step in the workflow. First, we test the Cilia.AI selections by running the pipeline through the “Deep Learning Segmenter”. This allows users to see how well it works on the given dataset and adjust parameters as needed. Second, for each pipeline, there are a series of specific parameters that the user must adjust as needed based on their staining and intended analysis. These parameters should be checked for each independent dataset to ensure that the resulting selection is as desired **(Video S5, S6, S7)**. When changes are made to any pipeline, the pipeline must be exported in order to be run with those changes. To analyze large datasets, we recommend operating the SubQ pipelines in the batch analysis feature of the ZEISS arivis Pro software. Before running a batch analysis, the MIP images should be imported into the ZEISS arivis Pro software, where they are automatically converted and saved into a single SIS file **(Video S8)**. The primed pipeline and SIS file can then be used to carry out batch analysis (**Figure S5C).**

To export data for analysis, each pipeline has a distinct export module with specific output tags. While it is possible to export individual data such as sums, average intensities, and other features per image, this can be quite time-consuming. To expedite the export process, we developed three individual data export pipelines that are tailored to each specific SubQ pipeline. We decided to keep data export separate from the detection pipelines to allow for manual corrections. Each data export pipeline imports the stored document objects with the corresponding tags predefined by its respective main SubQ pipeline **(File names: SubQ_BB_DC Batch Export; SubQ_CT Batch Export; SubQ_TZ Batch Export)**. It then uses an export function in which features can be selected and analyzed for any desired subcompartments **(Figure S5D)**. It is critical to ensure that the export function stores the exported data in the same directory as the SIS file. Once the features and directory are chosen, the export pipeline can be saved with these settings and used in batch analysis. The export directory has to be the same as the directory containing the SIS file, and the file names must be shortened **(Figure S5E)**. If the file names are not shortened, errors may occur that prevent data export due to an excessively long file path. After batch analysis for data export, there should be at least a single Excel spreadsheet per image in the SIS file **(Video S9)**.

### Leveraging Cilia.AI detection for cilia length analysis

To enhance the versatility of Cilia SubQ, we used our automated ciliary segmentation to measure cilia length, a key indicator of ciliary dysfunction that is often affected in cilia-related diseases. Cilia.AI is intrinsically embedded in each developed pipeline, allowing the extraction of cilia objects for subsequent analysis in FIJI. We therefore developed a separate export pipeline for use in conjunction with any of the SubQ pipelines to extract these ciliary objects. We recommend first using batch analysis to run the original SubQ pipeline and make any necessary manual corrections to cilia selections **(Figure S6A)**. Then, our SubQ_Length pipeline will import any stored objects with the tag “Primary cilia OUTPUT” and export them as a masked image **(Figure S6B)**. These must be exported to the same folder as the SIS file, or else the user may encounter errors when the pipeline is run **(Video S10)**. The pipeline is then saved with these changes and run as a batch analysis with the export directory matching the directory of the SIS file, and image names shortened **(Figure S6C, Video S10)**. This batch analysis produces one TIFF file per image, depicting a black-and-white selection of primary cilia.

For skeletonization and ciliary length analysis, we deployed a modified FIJI script (**Cilia Length_FIJI)**. Prior to running the FIJI script, all masked cilia images should be transferred to a separate empty folder to avoid errors when running the script. The script batch-processes the masked images and generates regions of interest (ROIs) and color-coded images with length measurements for each ROI **(Figure S6C, D, Video S10)**. The ROIs, color-coded images, and branch length measurements are automatically saved to the chosen output directory, which we recommend be an empty folder to avoid causing errors in the script. Saving the ROIs and color-coded images allows the ROIs to be overlaid on the original image if needed, and the measurements to be traced back to their corresponding color-coded cilia. For every image processed, there will be two Excel files with generated measurements. The first is a Cilia Length Statistics spreadsheet, which lists the cilia lengths. The second is a Skeleton Details spreadsheet, which provides information on any “branching” that was detected. We found that in certain cases, skeletonization of the primary cilia resulted in branched ROIs and therefore produced multiple length measurements, one for each branch. In these cases, there will be multiple measurements for the same cilia in the Cilia Length Statistics spreadsheet, which can either be combined to generate a single output or disregarded and manually measured instead.

To test the accuracy of the overall workflow, we measured cilia length manually and compared it with Cilia.AI-based quantifications in two cell lines, MEFs and human skin fibroblasts. We found that the automated method performed similarly to expert manual annotations, indicating reliable precision **(Figure S6E)**. To assess redundancy and flexibility across the three pipelines, we compared the performance of SubQ_BB_DC, SubQ_CT, and SubQ_TZ. All three detected a similar number of cilia per field of view and showed comparable performance in cilia length quantification, indicating that they are interchangeable and provide analytical flexibility **(Figure S6F)**.

### Kymograph generation for primary cilia trafficking

To further expand the SubQ suite, we utilized the ZEISS arivis Pro platform to analyze intrinsic trafficking within the primary cilium. Such dynamics are often used to decipher defective anterograde and retrograde trafficking as well as perturbations in entry and exit of ciliary molecules. The goal is to create a unified platform to generate kymographic outputs that facilitate quantification of molecular trafficking, especially using IFT machinery as a key readout. To examine IFT trafficking within the cilium, we transfected MEFs with IFT88-GFP and ARL13B-mCherry and recorded short videos at one image per 250 ms for 30 seconds **(Video S11 and S12)**. If a video contains multiple cilia, each cilium must be cropped into its own image using the Transformation Gallery under the Data Tab in the ZEISS arivis Pro software. To do this, we opened the video (.nd2 file, Nikon) and selected the crop tool to create individual videos for each cilium. Cropping should be tightly restricted to the cilium to minimize background signal. Each subsequent step must be performed on each cropped cilium video. Prior to kymograph generation, background correction may be applied to reduce noise, although in our experience this step can be inconsistent. In the Transformation Gallery (Data tab), we then rotated each cilium individually around the Z-axis so it lay horizontally, with the centrosome on the left and resampling set to bicubic interpolation. To generate a kymograph, we manually annotated the cilium on the first frame using the Draw Object tool (blue circle and pen). It is critical to avoid selecting non-ciliary background to ensure accuracy. A “Manual” object was next created, and the kymograph Python script **(SubQ_Kymo)** was then executed on the annotated video, generating an initial kymograph image **(Figure 5A, Video S13)**. The user will need to use the Brightness Auto Adjust function to visualize it properly. To further enhance kymograph visualization, we resampled the pixel size in the x– and y-axes, multiplying each by 10, and changed the pixel type from 16-bit to 32-bit float **(Figure 5B, Video S13)**. Each kymograph plot can be exported as a .tif file and the complete arivis project can then be saved as a .SIS file. Kymograph quantification can then be performed manually using FIJI software. SubQ_kymo is a semi-manual visualization workflow and a valuable add-on to the Cilia SubQ toolkit that can be expanded and further enhanced to streamline the manual analysis of ciliary events and their dynamics.

**Figure 5:**
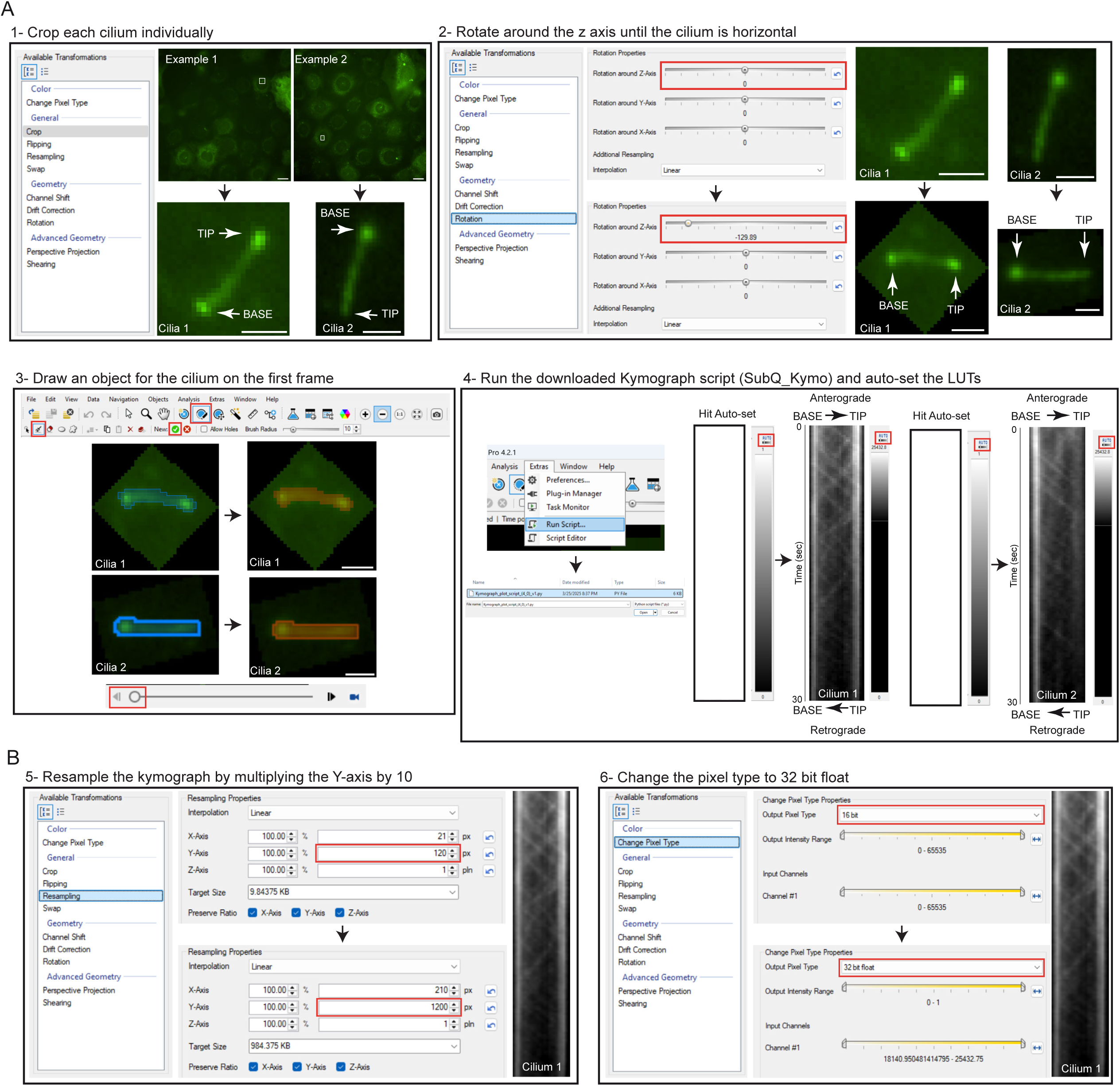
Generating kymographs in Zeiss Arivis Pro. **A.** Visual representation of how videos are processed in Arivis Pro software, including manual annotation of the primary cilium and execution of SubQ_Kymo script to generate the kymograph, along with two representative examples (scale bar in full-view images: 20 µm, scale bar in cropped and rotated images: 2 µm). **B.** Representative example depicting the post-processing steps applied to the kymograph to facilitate subsequent analysis.

## Discussion

Cilia SubQ is a collection of pipelines and scripts that researchers can adapt to their experimental needs. Our goals are to streamline time-consuming tasks and provide unbiased, fully traceable analysis of large datasets. Although Zeiss Arivis Pro Software is not open-source, we provide pipeline files, ready-to-use scripts, export modules, and associated videos for the workflows tested and validated in this manuscript. In addition, the underlying software is widely used and available in various academic institutions, making this suite deployable across these imaging environments. One of the key features that motivated our selection of Zeiss Arivis Pro software was its associated training platform, ZEISS arivis Cloud, which provides a stable and reliable environment for the consistent, continuous training of our Cilia.AI model. This platform supports iterative optimization while preserving previous training versions, thereby capturing ciliary diversity across experiments. The resulting model is fully supported within the software and can be integrated into any cilia-analysis pipeline. Preserving previous versions of Cilia.AI was also critical as a safeguard against performance loss following additional rounds of training.

### Performance, scope, and limitations of Cilia.AI

The Cilia.AI module is the core ciliary detector within the suite and performs robustly across multiple cell types and imaging conditions. Cilia.AI was thoroughly tested, and its performance was evaluated on previously unseen datasets from three different cell lines. The similar behavior observed across training and non-training datasets suggests that the model performs adequately under our current imaging conditions and may have approached a practical performance plateau. Importantly, Cilia.AI not only reproduced manual object detection, but also captured biological changes in ciliary protein levels, as illustrated by the SAG-dependent reduction in GPR161 signal. Although the suite was primarily developed and validated in monolayer cell systems, the additional proof-of-concept testing in mature mouse brain sections suggests that our cilia annotator retains utility in more complex in vivo contexts, albeit with lower performance and greater reliance on manual correction. In this context, Cilia.AI performance remains dependent on staining quality, signal-to-noise ratio, and imaging conditions, all of which can lead to occasional missed or partially selected cilia. In rare instances, Cilia.AI may classify non-ciliary fluorescent signal or excised ciliary tips as primary cilia, although in our hands these events were uncommon and typically affected fewer than 3% of objects. This underscores the need for expert oversight to identify and remove any unwanted objects when present.

We found that Cilia.AI performance varied depending on the ciliary marker and reagent used. For example, a mouse ARL13B antibody provided ciliary labeling with minimal non-ciliary background, yielding the most reliable detection. In contrast, a rabbit ARL13B antibody (Proteintech, 17711-1-AP) produced higher non-ciliary background and reduced performance. Other ciliary markers not explicitly tested in this study, such as CFAP300^56^, may be compatible with Cilia.AI, provided they generate predominantly ciliary signal with minimal background. In addition, Cilia.AI sensitivity and specificity can be hindered by poor staining or suboptimal imaging quality, including very low ARL13B intensity, defined here as below 30 A.U. in 8-bit maximum-intensity-projection images. This underscores the requirement for medium-to-high, non-saturated ARL13B fluorescent signal for robust segmentation. For these reasons, expert visual confirmation remains an essential part of the workflow, even though the user’s role shifts from labor-intensive annotation toward a more supervisory review.

### Comparison of Cilia SubQ to existing tools

In recent years, the cilia field has developed key resources that automate the detection and analysis of primary cilia to reduce the time, bias, and variability associated with manual ciliary annotation. Among the relevant tools are CiliaQ (Java), detectCilia (R), ACDC (MATLAB), and AI-based workflows implemented in NIS Elements (Nikon), each of which addresses the analysis of primary cilia using different programming languages and software environments^39–42^. CiliaQ occupies a broad analytical space and is an open-source FIJI-based workflow that supports 2D, 3D, and 4D datasets from cell lines and tissues. It provides a variety of outputs, including ciliary length, bending, orientation, and fluorescence intensity, and supports batch analysis with proposed segmentation methods and analysis of fluorescence distributions along the cilium^40,43^.

While CiliaQ can generate a wide range of full-cilium outputs, Cilia SubQ offers a broad collection of pipelines and scripts that spans analysis from standard 2D full-cilium parameters to quantifying both centrioles and ciliary subdomains, complemented by the generation of kymographic plots. While there is some overlap between the two methods in outputs, including cilia length and fluorescence intensity, the central design of Cilia SubQ is different: it employs cilium detection to segment ciliary subdomains, including the basal body, daughter centriole, transition zone, and ciliary tip. Cilia SubQ and CiliaQ are intrinsically different but complementary in purpose and use. Cilia SubQ also offers analytical traceability by using a single SIS file that can store pipeline parameters, images, annotations, and corrections. This is especially important when working with challenging subdomains or subtle phenotypes, without disrupting the entire workflow.

### Centrosome and ciliary subdomain analysis

The SubQ_BB_DC pipeline extends Cilia.AI-based cilium detection to the centrosome, enabling basal body and daughter centriole segmentation. In MEFs, the pipeline showed strong performance because centrioles were sufficiently separated. In contrast, RPE-1 cells and skin fibroblasts exhibited more closely apposed centrioles with γ-tubulin labeling, reducing detection accuracy and requiring manual correction. In this context, a more defined core centriolar marker such as Centrin-2 can improve centriole delineation. By refining blob-finder parameters, particularly object diameter and probability thresholds, the SubQ_BB_DC pipeline can be adapted to reliably capture centrosomal signal and distinguish between the basal body and the daughter centriole.

The same logic applies to other centrosomal markers not explicitly tested here. For proteins with spatial distributions similar to γ-tubulin or Centrin-2, existing parameters can be reused as a starting point, followed by local refinement. The ZEISS Arivis Pro can uniformly dilate or shrink pre-existing basal body or daughter centriole selections with a specified number of pixels. This allows users to adapt segmentation masks to proteins with broader or narrower spatial distributions while preserving consistent projected areas across conditions. The user can also add a “Segment Morphology” step, with either “Basal body OUTPUT” or “Daughter centriole OUTPUT” as the input. This approach is sufficiently flexible to accommodate additional centrosomal markers with the necessary adjustments, testing, and validation.

Similarly, the SubQ_TZ and SubQ_CT pipelines provide a framework for quantifying the transition zone and ciliary tip. These pipelines can be adapted to detect other proteins of interest. To this end, when tuning blob finder parameters, we recommend balancing probability threshold and selected diameter to capture all transition zone– and ciliary tip-positive signals, including non-ciliary areas. Our design accounted for this by implementing layers of stringency in both pipelines to filter out non-ciliary noise. For both subdomains, the user can also choose standardized markers as scaffolds to define compartments at widefield resolution and quantify other colocalizing markers, particularly those difficult to segment. If the reference marker signal does not fully encompass the region of interest, selections can be expanded or shrunk as needed to match the exact localization of the protein of interest.

### Manual correction strategies and use cases

Cilia SubQ pipelines are designed for targeted and efficient manual intervention. For primary cilia, manual corrections are necessary only when measuring the entire cilium, such as cilium length or average intensity of a ciliary marker. In these cases, users should delete or retrace partially detected or undetected cilia, and newly created objects must receive the appropriate output tag. On the other hand, if users want to focus only on specific subcompartments, such as the transition zone, tip region, or basal body, they can limit corrections to the relevant subdomain. This focused approach reduces hands-on time while maintaining the accuracy of measurements within the area of interest.

### Key limitations of this study

Cilia SubQ was mainly tested in monolayer cells and mouse brain sections, but it was not broadly extended to other models, tissues, or organoids. This study focused on assessing ciliary subdomains. We used 2D MIP images for proof-of-concept validation and did not include 3D or time-resolved imaging during pipeline development and testing. As a result, this study does not confirm the performance of Cilia SubQ for 3D or 4D datasets. Additional optimization and validation will be needed before the framework can be confidently applied in those areas. KymographClear and KymographDirect are specialized, publicly available tools for creating and analyzing kymographs^57^. Our kymograph script integrates into ZEISS arivis Pro software, offering users a more unified cilia-analysis environment. However, our kymograph generation workflow does not include integrated measurement of speed or event frequency for IFT trafficking. Furthermore, our attempts to automate IFT event tracing gave inconsistent results, indicating that this part is not yet reliable for quantitative analysis.

## Supporting information

Cilia SubQ files

Cilia S12 Trafficking Example 2

Video S1 – SubQ_BB_DC

Video S2 – SubQ_TZ

Video S3 – SubQ_CT

Video S4 – Prime the pipeline

Video S5 – SubQ_BB_DC changes to pipeline

Video S6 – Sub_TZ changes to pipeline

Video S7 – SubQ_CT changes to pipeline

Video S8 – Make a SIS file

Video S9 – Data Export

Video S10 – SubQ_Length (+FIJI)

Video S11 Cilia Trafficking Example 1

Video S13 – SubQ_Kymo

## Supplementary figures

**Supplementary Figure 1:** Validation of Cilia.AI on images used to train Cilia.AI and on brain tissue images. Related to Figure 1. **A.** Graphs showing the cilia detection performance of Cilia.AI in MEFs, RPE-1 cells, and human skin fibroblasts on datasets used to train Cilia.AI. For each cell line, the first graph shows the percentage of cilia detected per image (one-sample Wilcoxon signed-rank test); and the second graph displays the number of cilia detected per image (Friedman’s test). **B.** Primary cilia in P31 cortical and hippocampal mouse brain tissue sections, with and without Cilia.AI detection (scale bar: 10 µm). **C.** Graphs displaying the cilia detection performance of Cilia.AI in P31 mouse brain tissue sections unseen during Cilia.AI training. The left graph shows the percentage of cilia detected per image in mouse brain tissue, analyzed with a one-sample Wilcoxon signed-rank test. The graph on the right displays the number of cilia detected per image in mouse brain tissue, analyzed with Friedman’s test. **D.** Statistics comparing manual and Cilia.AI cilia annotation. Manual statistics are estimates based on the ratio calculated from manually annotating 75 primary cilia. For panel A, a total of 6 independent experiments (1009 primary cilia) were analyzed, with two experiments per cell line. For panel C, a total of 729 cilia were quantified in mouse brain tissue sections.

**Supplementary Figure 2:** Detection of basal bodies labeled with γ-tubulin in RPE-1 and human skin fibroblasts. Related to Figure 2. **A.** Graphs showing the basal body detection performance of SubQ_BB_DC in RPE-1 cells stained for γ-tubulin (centrosome marker). The left graph shows the percentage of basal bodies correctly detected per image, analyzed with a one-sample Wilcoxon signed-rank test. The middle graph shows the number of basal bodies detected per image, analyzed using paired Wilcoxon signed-rank test. The right graph shows the average γ-tubulin intensity of basal bodies in RPE-1 cells detected either manually (n=75 cells) or with SubQ_BB_DC (n=489 cells). Statistical analysis was performed using an unpaired t-test. **B.** Same as A, but for human skin fibroblasts. Manual selections (n=75 cells) and SubQ_BB_DC annotations (n=230 cells). For panels A and B, a total of 4 independent experiments were analyzed, with two per cell line, and a total of 857 primary cilia were quantified across both cell lines.

**Supplementary Figure 3:** Challenges of apposed centrioles and validation of SubQ_BB_DC in annotating the daughter centriole. Related to Figure 2. **A.** Illustration of how two closely apposed centrioles can be detected by SubQ_BB_DC as a single object rather than two separate objects (scale bar: 2 µm). **B.** Depiction of incorrect centriole detection by SubQ_BB_DC when using a smaller diameter and a higher probability threshold (scale bar: 2 µm). **C.** The pipeline steps used to detect and isolate the daughter centriole in MEFs labeled with γ-tubulin in red and ARL13B in green. Cilia, basal body and daughter centriole annotations are shown as colored selections (scale bar: 2 µm). **D.** Graphs displaying daughter centriole detection performance of SubQ_BB_DC in MEFs stained for γ-tubulin. The top graph shows the percentage of daughter centrioles correctly detected per image, analyzed with a one-sample Wilcoxon signed-rank test. The bottom graph shows the number of daughter centrioles correctly detected per image, analyzed using paired Wilcoxon signed-rank test. **E.** Same as D, but for RPE-1 cells stained for Centrin-2. For panels D and E, a total of four independent experiments (823 primary cilia total) were analyzed.

**Supplementary Figure 4:** Rationale for layered filters in SubQ_TZ. Related to Figure 3. **A.** Depiction of multiple AHI1 transition zone signals associated with primary cilia in human skin fibroblasts (scale bar: 2 µm). **B.** A representative example in human skin fibroblasts stained with AHI1 showing that the correct transition zone signal is closer to the basal body than the non-specific AHI1 signal (scale bar: 2 µm). **C.** Display of the final AHI1 transition zone selection after application of the layered filters in SubQ_TZ (scale bar: 2 µm). **D.** Statistics comparing manual and SubQ_TZ transition zone annotation. Manual statistics are estimated from a timed annotation of five transition zone signals. SubQ_TZ correction statistics are estimated from correcting a single image.

**Supplementary Figure 5:** How to prime the pipeline and perform a batch analysis. Related to Figures 1, 2, 3, and 4. **A.** Visualization of the Deep Learning Segmenter operation, which imports and deploys the Cilia.AI model. Representative images displaying human skin fibroblasts labeled with ARL13B (green), γ-tubulin (red) and DAPI. Cilia annotations are shown as colored selections (scale bar: 10 µm). **B.** Demonstration of how to export a pipeline to save any changes. **C.** Overview of the steps required to run multiple images through a single pipeline using batch analysis. **D.** Illustration of the steps required to adjust the data export pipeline to obtain the desired quantifications. **E.** Images showing how to export quantifications when the data export pipeline is used in batch analysis.

**Supplementary figure 6:** Workflow for cilia length measurements. Related to Figure 1. **A.** Illustration of primary cilia detected by Cilia.AI in human skin fibroblasts. Cells are labeled with ARL13B (green), γ-tubulin (red) and DAPI. Cilia annotations are shown as colored selections (scale bar: 10 µm). **B.** Steps depicting how to export masked images of Cilia.AI cilia selections for batch analysis. **C.** A representative example of the exported masked image containing Cilia.AI selections in white and the background in black. **D.** Demonstration of the steps in the Cilia Length FIJI script used to obtain cilia length measurements. Cilia are first skeletonized and color-coded. ROIs are then drawn and assigned numbers based on the color-coding. **E.** Graphs showing cilia length measured either manually or with Cilia.AI+FIJI in MEFs and human skin fibroblasts. Both graphs were analyzed with unpaired t-tests. **F.** Graphs showing the detection performance of all SubQ pipelines in RPE-1 cells. The left graph shows the percentage of primary cilia detected per image, analyzed with a one-sample Wilcoxon signed-rank test. The middle graph shows the number of cilia detected per image, analyzed using Friedman’s test. The right graph shows the cilia length of detected primary cilia, analyzed with a one-way ANOVA. For panels E and F, a total of six independent experiments were analyzed, two per cell line. For the percentage of cilia detected per image and the number of cilia detected per image graphs panel in F, a total of 1,443 primary cilia were quantified across all three pipelines (481 primary cilia each).

**Video S1 – SubQ_BB_DC. Related to** Figure 2 – A video showing users how to import and run SubQ_BB_DC as a batch analysis, and what the results should look like. Note: the pipeline used in the video is the equivalent of SubQ_BB_DC. Additionally, the output “Daughter centrosome OUTPUT” is the equivalent of “Daughter centriole OUTPUT”.

**Video S2 – SubQ_TZ. Related to** Figure 3 – A video showing how to import and run SubQ_TZ as a batch analysis, and expected end state.

**Video S3 – SubQ_CT. Related to** Figure 4 – A video demonstrating how to import and execute SubQ_CT as a batch analysis, and what the results are expected to look like.

**Video S4 – Prime the pipeline. Related to** Figures 1**, 2, 3, and 4** – This video shows how users should prime their pipeline with the Cilia.AI model and export the edited pipeline prior to running any SubQ pipeline as a batch analysis. Note: The pipeline used in the video is the equivalent of SubQ_BB_DC. The cilia detection model imported into the deep learning segmenter in the video is the equivalent of Cilia.AI v32.

**Video S5 – SubQ_BB_DC changes to pipeline. Related to** Figure 2 – A video that highlights the steps in SubQ_BB_DC where the user is encouraged to make changes based on their specific staining. Note: The pipeline shown in the video is the equivalent of SubQ_BB_DC.

**Video S6 – Sub_TZ changes to pipeline. Related to** Figure 3 – A video showing the steps in Sub_TZ where the user is encouraged to make changes based on their specific staining.

**Video S7 – Sub_CT changes to pipeline. Related to** Figure 4 – A video showing the steps in SubQ_CT where the user is encouraged to make changes based on their specific staining.

**Video S8 – Make a SIS file. Related to** Figures 1**, 2, 3, and 4** – This video demonstrates how MIP images are imported into Arivis Pro and converted to a .sis file.

**Video S9 – Data Export. Related to** Figures 1**, 2, 3, and 4** – A video that depicts how to extract data to use for quantification after a SubQ pipeline has been run for detection.

**Video S10 – Cilia length. Related to** Figure 1 – This video shows how users deploy SubQ pipelines in Arivis Pro, followed by a cilia length script run in FIJI, to generate data for cilia length quantification. Note: The first pipeline used in the video is the equivalent of SubQ_BB_DC. The second pipeline used in the video is the equivalent of SubQ_Length. The Cilia Length Macro used in the video in FIJI is the equivalent of Cilia Length_FIJI.

**Video S11 – Cilia Trafficking Example 1. Related to** Figure 5 – Live imaging of a cell transfected with IFT88-GFP, in which trafficking of IFT cargoes is visible.

**Video S12 – Cilia Trafficking Example 2. Related to** Figure 5 – Live imaging of a cell transfected with IFT88-GFP, in which trafficking of IFT cargoes is visible.

**Video S13 – SubQ_Kymo. Related to** Figure 5 **-** A video showing how to generate a kymograph in Arivis Pro. Note: The script used to generate the kymograph in the video is the equivalent of SubQ_Kymo.

## Acknowledgements

We thank Kelly Graber and the Histology and Imaging Core at Sanford Research for their invaluable assistance. This core is funded by the NIH COBRE grant 1P30GM145398. We also thank the Sarah Goetz laboratory, where some of the data used in this manuscript were acquired. We thank the Zeiss Arivis support team for their assistance with this study, specifically Argaja Deepak Shende for generating the batch MIP script and Mariia Burdyniuk for producing the kymograph script. We also thank Dr. Sampath Rangasamy and Dr. Vinodh Narayanan for sharing the human skin fibroblasts. This work was supported by the National Institute of General Medical Sciences (award R35GM151229).

## Author Contributions

E.M. generated, optimized, and validated all pipelines. Data was generated and analyzed by E.M., K.H., and G.H. Research design, data analysis, and manuscript writing were carried out by E.M. and A.L. All authors read, reviewed and approved the final manuscript.

## Declaration of Competing Interest

The authors declare no competing financial interests

## Declaration of generative AI and AI-assisted technologies in the writing process

During the preparation of this work the author(s) provided Perplexity with a finalized language in order to fix typos and minimally improve the readability. After using this tool/service, the author(s) reviewed and edited the content and take(s) full responsibility for the content of the published article.

---

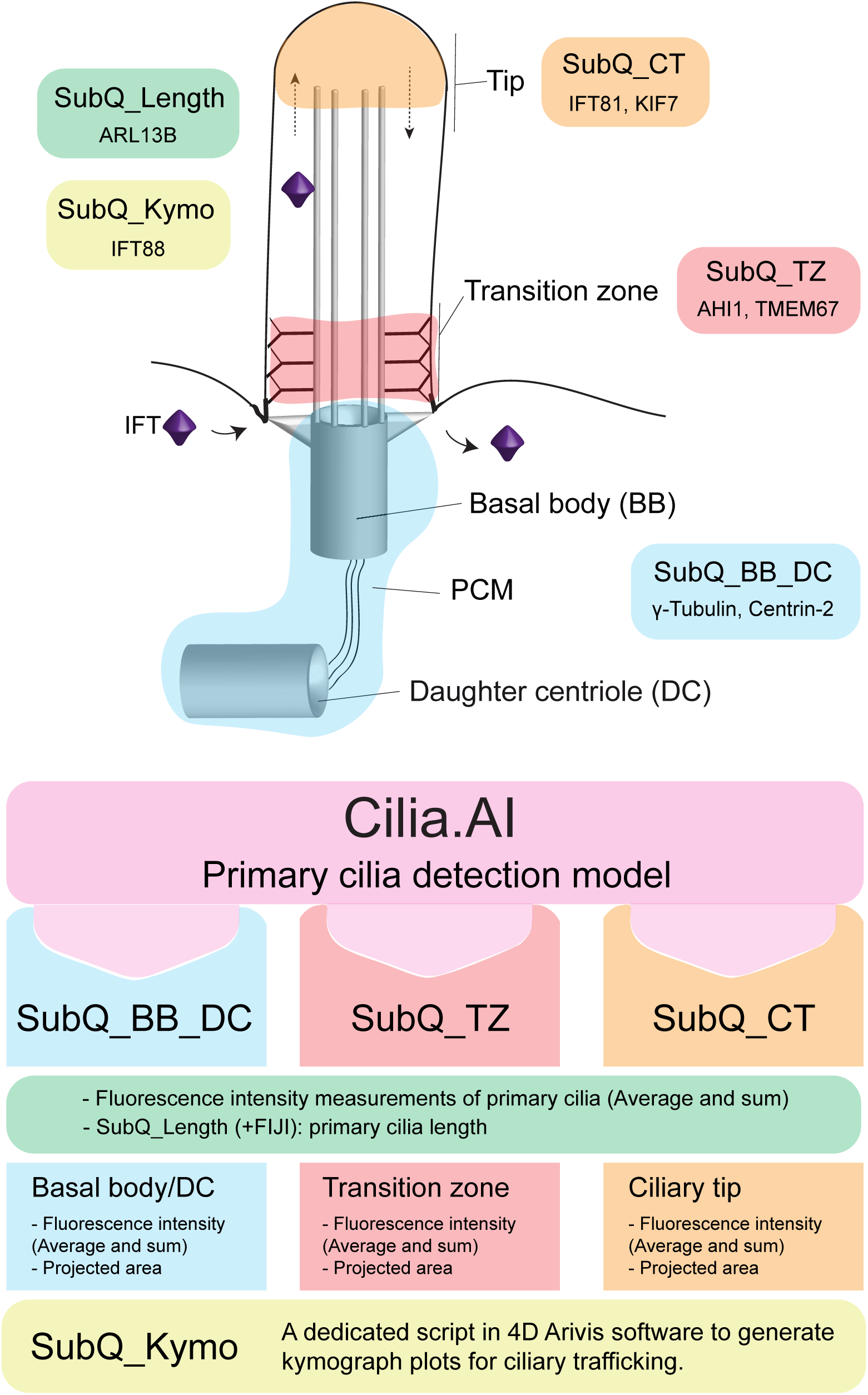
Graphical summary.

## Notes

https://osf.io/hm38f/

### Competing Interest Statement

The authors have declared no competing interest.

