## Supplementary material for "Cilia SubQ, a modular suite of pipelines for automated analysis of primary cilia and ciliary subdomains": Figure S1

A

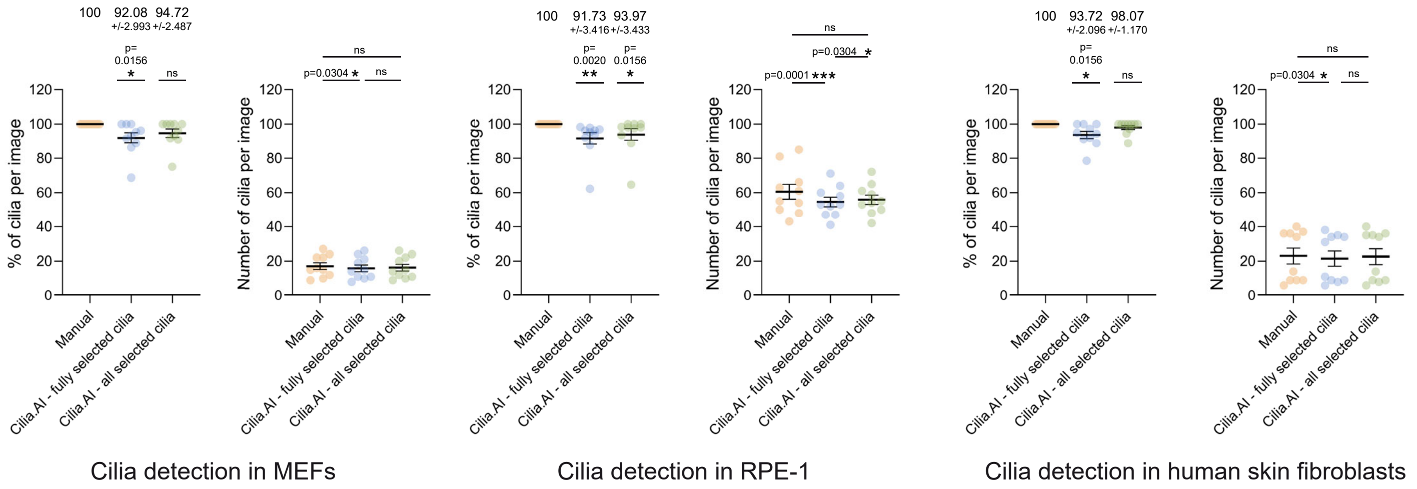


B

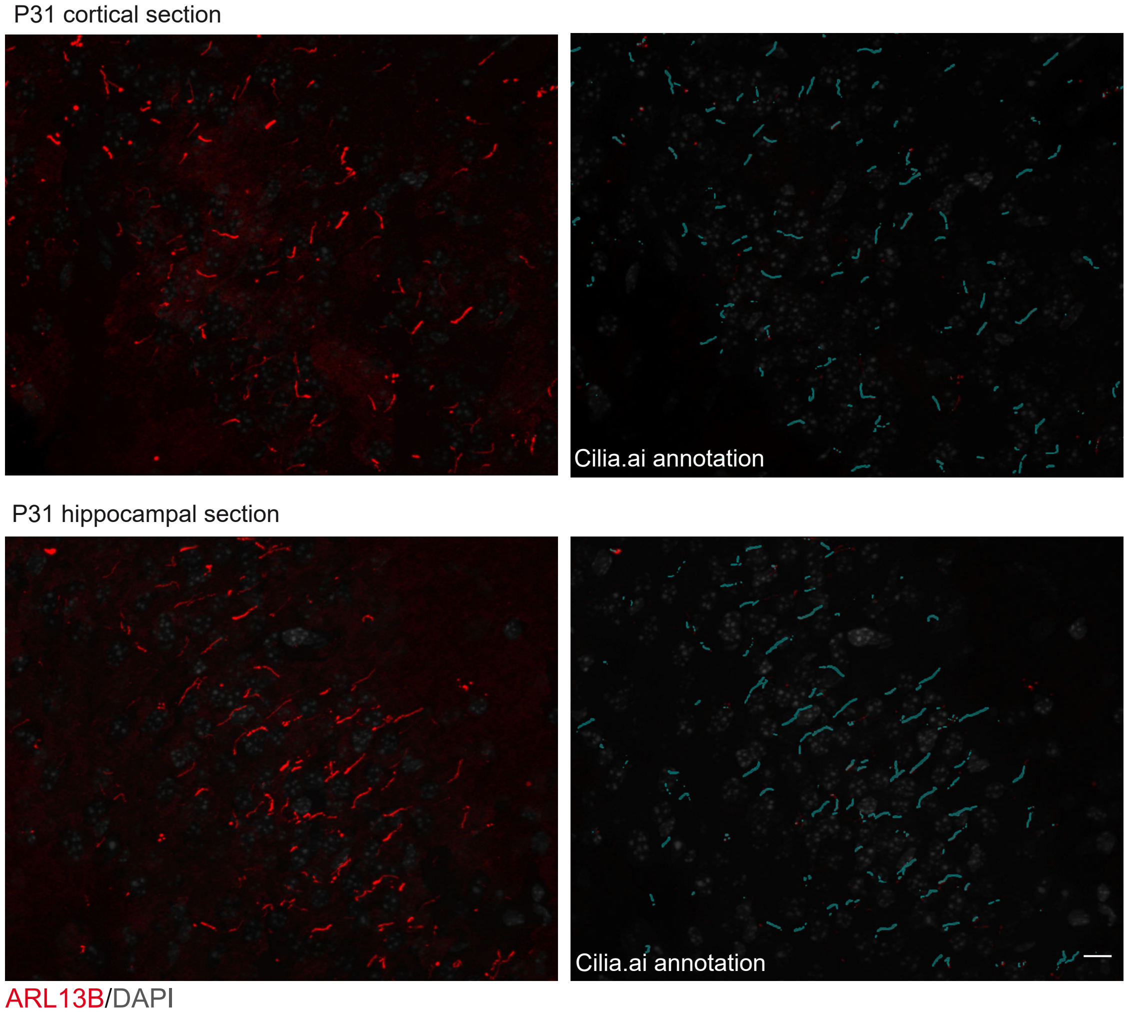


C

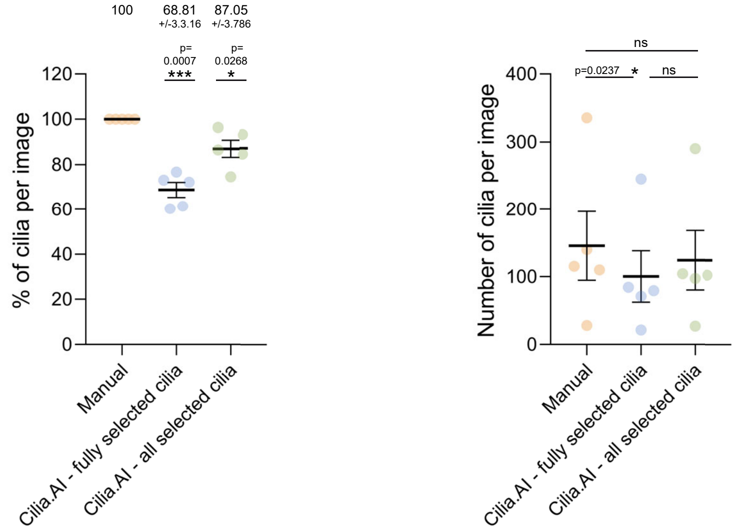


D

| | Manual | Cilia.AI |
|---|---|---|
| Number of images | 15 | 15 |
| Number of primary cilia | 234* | 234 |
| Processing time | 4 hr 10 min* | 10 min |
| Manual correction time | - | 20 min |
| Export data time | 1 hr 30 min* | 15 min |
| Total time | 5 hr 40 min* | 45 min |
| Fold change | - | 7.5X |
