## Supplementary figures and images for "Cilia SubQ, a modular suite of pipelines for automated analysis of primary cilia and ciliary subdomains"

### Figure S2

Figure S2

A

Basal Body-PCM detection in RPE-1

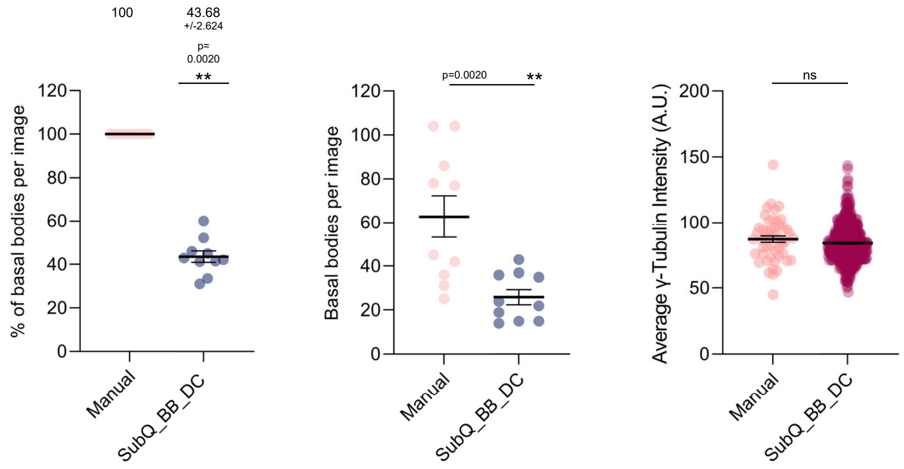


B

Basal Body-PCM detection in skin fibroblats

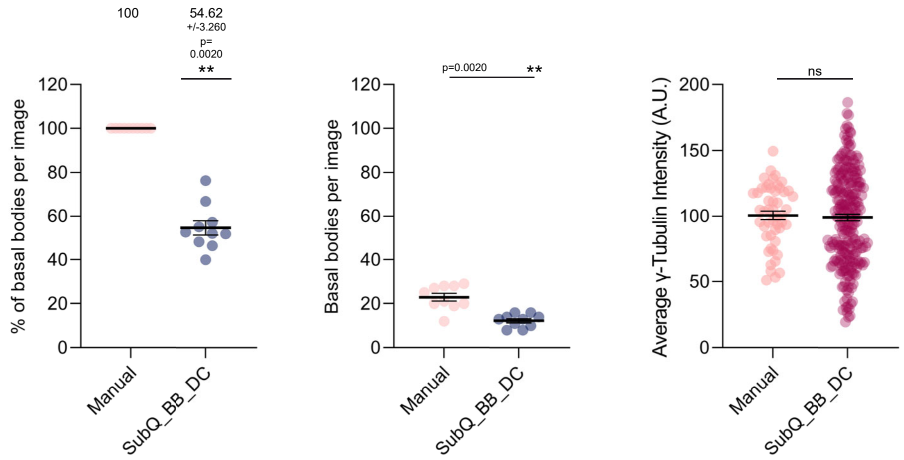

### Figure S3

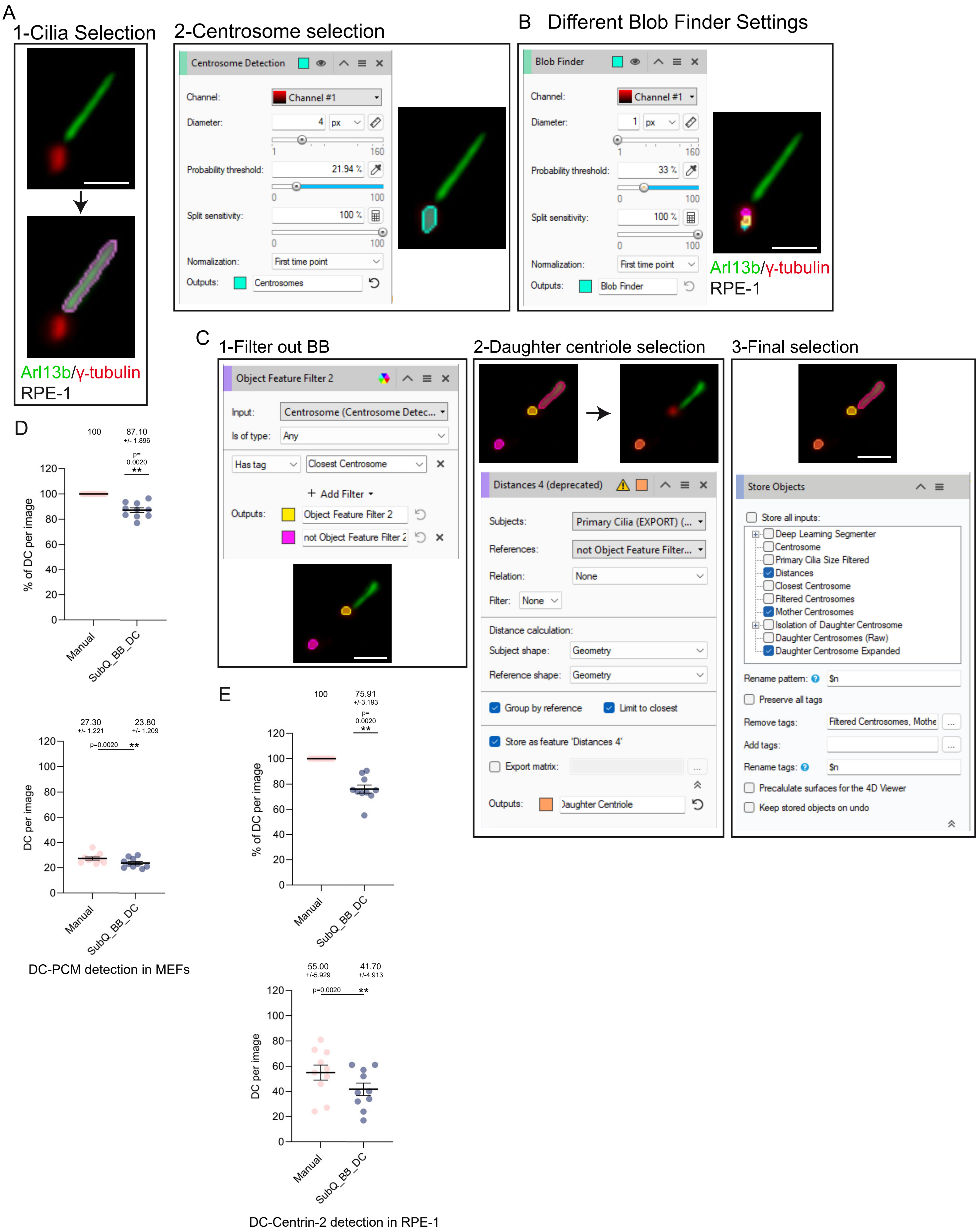

### Figure S5

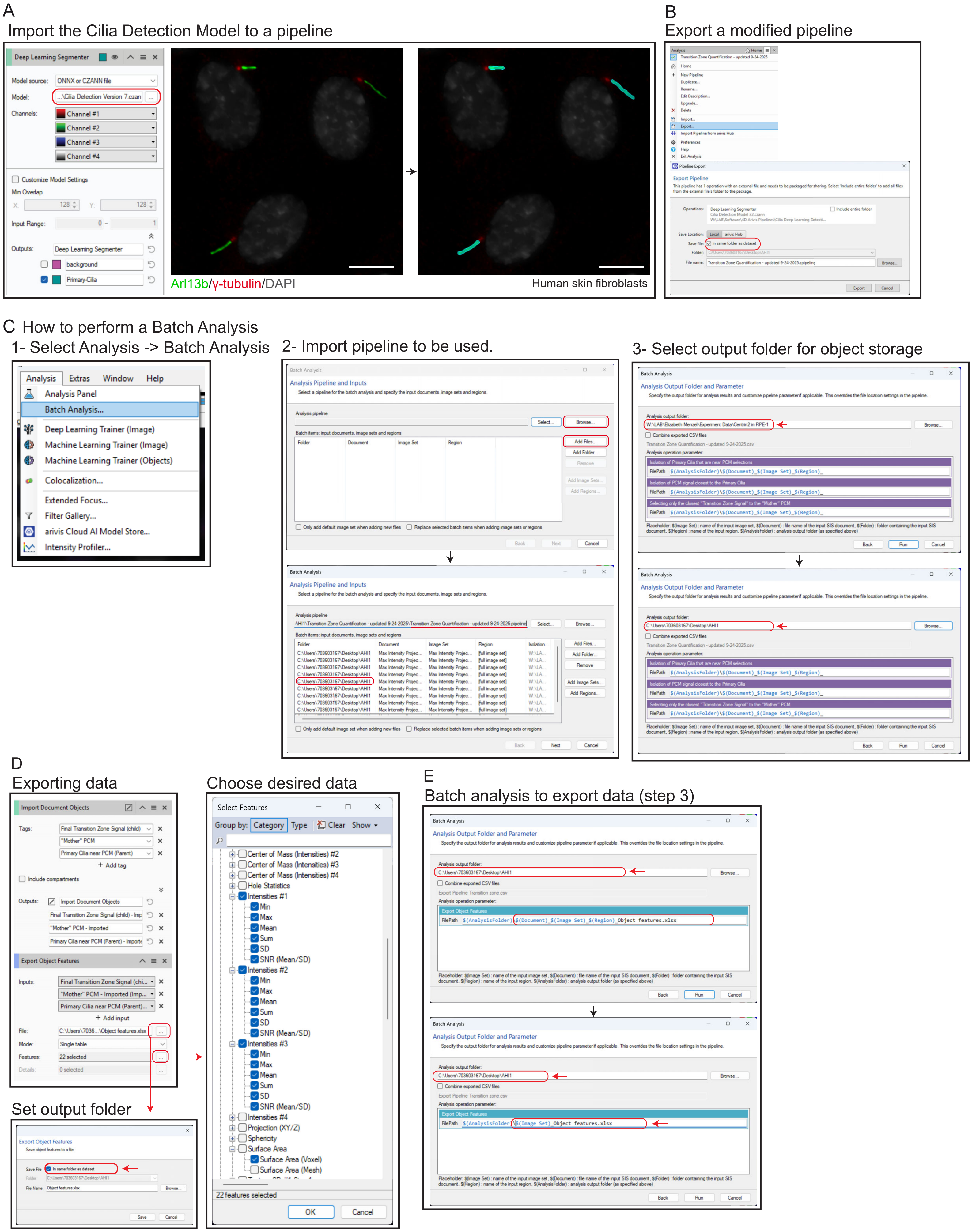

### Figure S6

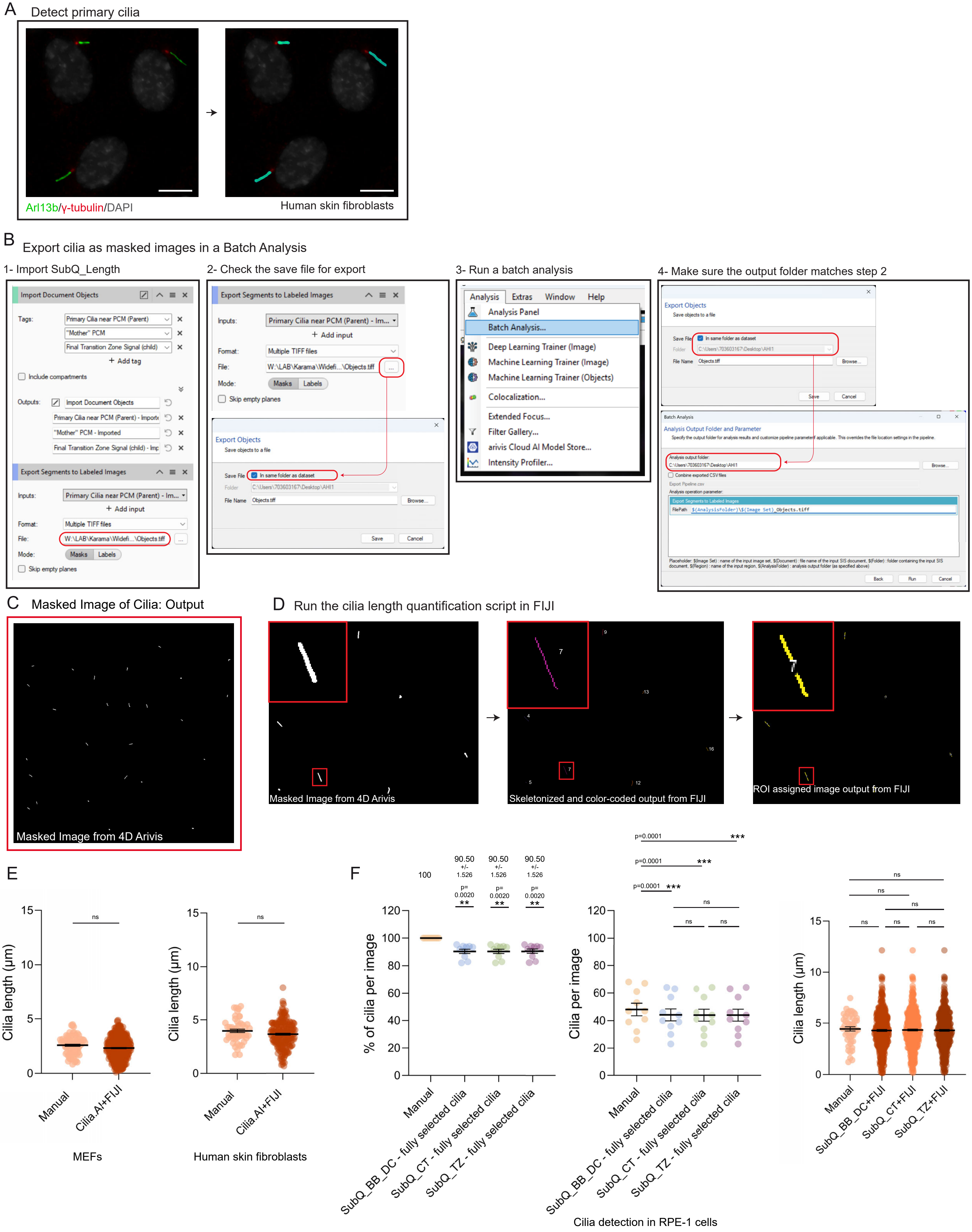
