## Supplementary material for "Cilia SubQ, a modular suite of pipelines for automated analysis of primary cilia and ciliary subdomains": Figure S4

A

Step 5 (Fig 3B) selection

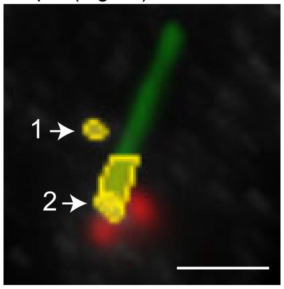


AHI1/Arl13b/
γ-tubulin/DAPI
Human skin fibroblasts

B

Step 6 (Fig 3B) selection

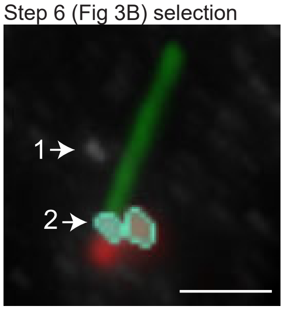


C

Final selection

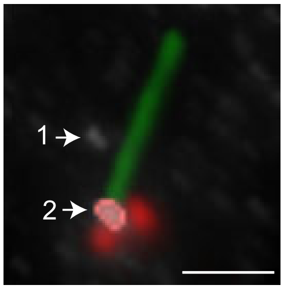


D

| | Manual | SubQ |
|---|---|---|
| Number of images | 5 | 5 |
| Number of primary cilia | 102* | 102 |
| Processing time | 2 hr 30 min | 2 min |
| Manual correction time | - | 13 min |
| Export data time | 1 hr 30 min | 15 min |
| Total time | 4 hr | 30 min |
| Fold change | - | 8X |
