## Supplementary material for "Cilia SubQ, a modular suite of pipelines for automated analysis of primary cilia and ciliary subdomains": Cilia SubQ files: Cilia Length_FIJI.ijm

```
inputDir=getDirectory("Choose directory containing input files");
fileList=getFileList(inputDir);
outputDir=getDirectory("Choose directory to save files to");
run("Bio-Formats Macro Extensions");
for(q = 0; q <= fileList.length-1; q++) {
	print(fileList[q]);
	id = inputDir+fileList[q];
	Ext.setId(id);
	Ext.getSeriesCount(seriesCount);
	Ext.getSeriesName(seriesName);
	print(seriesName);
	print("opening "+fileList[q]);
	open(inputDir+fileList[q]);
	//opens the raw file
	filename = fileList[q];
	no_tiff = filename.replace(".tiff", "");
	fns = no_tiff.replace(" ","");
	print(fns);
	id2=getImageID();
	print("ID2: "+id2);
	setAutoThreshold("Default dark");
	run("Threshold...");
	setThreshold(255, 255);
	setOption("BlackBackground", true);
	run("Convert to Mask");
	run("Skeletonize");
	run("Analyze Skeleton (2D/3D)", "prune=none show display");
	selectWindow("Tagged skeleton");
	close();
	//closes the tagged skeleton image that is automatically generated with Analyze Skeleton
	selectWindow(fns+"-labeled-skeletons");
	id3 = getImageID();
	print("ID3: "+id3);
	save(outputDir+no_tiff+" Color-coded.tiff");
	//id3 is the image with the color-coded skeleton
	run("Select All");
	getStatistics(area, mean, min, max, std, histogram);
	for (i = 1; i < max+1; i++) {
		setThreshold(i-0.5, i+0.5);
		run("Create Selection");
		roiManager("Add");
	};
	selectWindow("ROI Manager");
	roiManager("Show All with labels");
	selectWindow("ROI Manager");
	roiManager("Save", outputDir+no_tiff+" Cilia Length ROIs.zip");
	wait(3000);
	selectImage(id3);
	selectWindow("ROI Manager");
	run("Flatten");
	id4 = getImageID();
	print("ID4: "+id4);
	//id4 is the flattened image
	selectImage(id4);
	save(outputDir+no_tiff+" Cilia Length Flattened.tiff");
	selectWindow("Branch information");
	saveAs("Results", outputDir+no_tiff+" Cilia Length Statistics.csv");
	selectWindow(no_tiff+" Cilia Length Statistics.csv");
	close(no_tiff+" Cilia Length Statistics.csv");
	selectWindow("Results");
	saveAs("Results", outputDir+no_tiff+" Skeleton Details.csv");
	selectWindow("Results");
	close("Results");
	selectWindow("Threshold");
	close("Threshold");
	print("closed");
	selectImage(id2);
	setOption("Changes", false);
	close();
	roiManager("Deselect");
	roiManager("Delete");
};
print("Finished");

	
	
	
	
```
