## Supplementary material for "Cilia SubQ, a modular suite of pipelines for automated analysis of primary cilia and ciliary subdomains": Cilia SubQ files: MIP_8-bit conversion.ijm

```
inputDir=getDirectory("Choose directory containing input files");
fileList=getFileList(inputDir);
outputDir=getDirectory("Choose directory to save files to");
run("Bio-Formats Macro Extensions");
//setBatchMode(true);

for(i = 0; i <= fileList.length-1; i++) {
	print(fileList[i]);
	id = inputDir+fileList[i];
	Ext.setId(id);
	Ext.getSeriesCount(seriesCount);
	Ext.getSeriesName(seriesName);
	print(seriesName);
	print(seriesCount+" series in "+id);
	print("Series Count: "+seriesCount);
	for (k=0; k<seriesCount; k++) {
		print("opening "+fileList[i]+" series "+(k+1));
		run("Bio-Formats Importer","open=["+id+"] view_stack_with=Hyperstack series_"+(k+1));
		print("Field of view: "+(k+1));
		id2 =getImageID();
		print("Image ID: "+id2);
		run("Make Composite");
		run("Z Project...", "projection=[Max Intensity]");
		id3 = getImageID();
		selectImage(id3);
		print("New Image ID: "+id3);
		run("8-bit");
		run("Stack to Hyperstack...", "order=xyczt(default) channels=4 slices=1 frames=1 display=Composite");
		Stack.setChannel(1);
		run("Magenta");
		Stack.setChannel(2);
		run("Red");
		Stack.setChannel(3);
		run("Green");
		Stack.setChannel(4);
		run("Blue");
		id4=getImageID();
		print("Processed image ID: "+id4);
		print(fileList[i]+" series "+(k+1)+" converted to 8-bit");
		print(fileList[i]+" series "+(k+1)+" processed");
		selectImage(id2);
		setOption("Changes",false);
		close();
		selectImage(id4);
		title="Max Intensity Projection "+fileList[i]+" Z-stack "+(k+1);
		save(outputDir+title+".tif");
		print(title+" saved");
	}	
}
print("Finished");
```
