## Supplementary material for "Cilia SubQ, a modular suite of pipelines for automated analysis of primary cilia and ciliary subdomains": Cilia SubQ files: SubQ_Kymo.py

```
# coding: utf-8


import time
import arivis
import arivis_objects as objects
import arivis_core as core
import numpy as np

#%% User defined variables
# channel for the kimograph:
INPUT_CHANNEL = 0   # <---- Count start from 0 (Ch#1 == 0)

TAG_DESCRIPTOR = "Manual"  # <--- Segments tag 

#%% Function : GetEnviroment
def GetEnviroment():

  viewer = arivis.App.get_active_viewer()
  imageset = viewer.get_imageset()
  if imageset is None :             #  if None == imageset:
    print("No Image Set open") 
    return viewer,None  
  return viewer,imageset

#%% Function : GetDocument
def GetDocument(imageset):
  if imageset is None:
    print ("No Image Set open" )
    return None  
  document = imageset.get_document()

  if document is None:
    print ("No Document open" )
    return None
  return document

#%% Function : GetPixelTypes()
def GetPixelTypes(imageset):
  
  if imageset is None :
    print( "No Image Set open" )
    return None   
  pixelType = imageset.get_pixeltype()
 
  if pixelType == core.ImageSet.PIXELTYPE_USHORT:
    typeP = 'H'    
    #dt = np.ushort
    dt = np.dtype(np.ushort) 
    Coefficent = int(65535)   
  elif pixelType == core.ImageSet.PIXELTYPE_ULONG:
    typeP = 'L' 
    #dt = np.uint
    dt = np.dtype(np.uint)
    Coefficent = int(65535)
  elif pixelType == core.ImageSet.PIXELTYPE_FLOAT:
    typeP = 'f' 
    #dt = np.float32
    dt = np.dtype(np.float32)
    Coefficent = int(65535)
  else:     #core.ImageSet.PIXELTYPE_UCHAR:
    Coefficent = int(255)
    typeP = 'B'
    #dt = np.uint8
    dt = np.dtype(np.uint8)
  return dt,typeP,Coefficent 

#%% Main

# helper to get execution time
startTime = time.time()
print ("Script is running ........ " )
#
viewer,imageset = GetEnviroment()
if imageset is not None:
    document = GetDocument(imageset)

    Store = document.get_store(imageset, objects.Store.DOCUMENT_STORE)
    channels = imageset.get_channel_count()
    timepoint_count = imageset.get_timepoint_count()
    _,typeP, _  = GetPixelTypes(imageset)
    
    if INPUT_CHANNEL < channels:    #if input channel exists
      print ("# Channels: " +  str(channels) + "  Input channel #: " + str(INPUT_CHANNEL) )
      
      if Store is not None :          # Get the storage
        IdList = Store.get_object_ids(TAG_DESCRIPTOR)
        
        if IdList.__len__() == 0:
          print( "No object with the specified tag exist. End of script!" )
        if IdList.__len__() == 1:
          Object = Store.get_object(IdList[0], True)
          print( "ID: " + str(IdList[0]) + "  # TP: " + str(Object.get_timepoint()))
          
        if IdList.__len__() >= 2:
          print( "More then one object with the specified tag exist. End of script!" )
        
        # get the objects bounds to prepare the bufer
        objBounds4D = Object.get_bounds()   # 4D bounds
        objBounds2D = core.Bounds2D()       # initianlizing 2D bounds
        objBounds2D.x = objBounds4D.x1
        objBounds2D.y = objBounds4D.y1  
        objBounds2D.height = objBounds4D.y2 - objBounds4D.y1 + 1
        objBounds2D.width  = objBounds4D.x2 - objBounds4D.x1 + 1
        
        obj_mask = np.zeros((objBounds2D.height, objBounds2D.width), dtype = bool)
        Object.get_mask(obj_mask, 0, objBounds2D) 

        obj_buffer = np.zeros((objBounds2D.height, objBounds2D.width), dtype = int )
        obj_buffer_T = np.zeros((objBounds2D.height, objBounds2D.width, timepoint_count), dtype = int) 
                    
        boundingBox3D = core.Bounds3D()
        boundingBox3D.x1 = objBounds4D.x1
        boundingBox3D.x2 = objBounds4D.x2
        boundingBox3D.y1 = objBounds4D.y1
        boundingBox3D.y2 = objBounds4D.y2
        boundingBox3D.z1 = 0
        boundingBox3D.z2 = 0
        
        line1D = np.zeros((timepoint_count, objBounds2D.width), dtype = float)
        for TP in range(0, timepoint_count):
            obj_buffer =  imageset.read_imagedata(region=boundingBox3D, channel=INPUT_CHANNEL, timepoint=TP) 
            obj_buffer[~obj_mask] = 0
            for i in range(0, objBounds2D.width):
                col = obj_buffer[:,i]
                col_mask = obj_mask[:,i]
                val =  np.mean(col[col_mask])
                line1D[TP,i] = val
#%%
        line1D_32bit = np.float32(line1D)
        
        get_bounding_box = imageset.get_bounding_rectangle(0)  # width height only
        get_bounding_box.width = np.shape(line1D)[1]  
        get_bounding_box.height = np.shape(line1D)[0]   
        get_bounding_box.x = 0
        get_bounding_box.y = 0
        
        DestBoundingBox3D = core.Bounds3D()
        DestBoundingBox3D.x1 = 0
        DestBoundingBox3D.x2 = np.shape(line1D)[1] - 1
        DestBoundingBox3D.y1 = 0
        DestBoundingBox3D.y2 = np.shape(line1D)[0] - 1
        DestBoundingBox3D.z1 = 0
        DestBoundingBox3D.z2 = 0
        
        DEST_IMAGESET_NAME = imageset.get_name() + '_rotated'
                      
        if imageset.get_id() == DEST_IMAGESET_NAME:
              raise RuntimeError("Already on the Destination dataset / Cannot delete current image set")               
        if document.get_imageset(DEST_IMAGESET_NAME) is not None:
              document.delete_imageset(document.get_imageset(DEST_IMAGESET_NAME))

        destImageset = document.create_imageset(DEST_IMAGESET_NAME, core.ImageSet.PIXELTYPE_FLOAT, 1)    # 16 bit  1=channels      
        destImageset.insert_timepoints(0, 1)            # timepoint 0 to sum of TPs (1)
        destImageset.insert_planes(0, 0, 1, get_bounding_box)  # timepoint first plane NfPlanes bounds
        destImageset.insert_channels(0, 0)

        destImageset.write_imagedata(line1D_32bit, DestBoundingBox3D, 0, 0)
```
